# Single-cell multi-omics of human clonal hematopoiesis reveals that *DNMT3A* R882 mutations perturb early progenitor states through selective hypomethylation

**DOI:** 10.1101/2022.01.14.476225

**Authors:** Anna S. Nam, Neville Dusaj, Franco Izzo, Rekha Murali, Robert M. Myers, Tarek Mouhieddine, Jesus Sotelo, Salima Benbarche, Michael Waarts, Federico Gaiti, Sabrin Tahri, Ross Levine, Omar Abdel-Wahab, Lucy A. Godley, Ronan Chaligne, Irene Ghobrial, Dan A. Landau

## Abstract

Somatic mutations in cancer genes have been ubiquitously detected in clonal expansions across healthy human tissue, including in clonal hematopoiesis. However, mutated and wildtype cells are morphologically and phenotypically similar, limiting the ability to link genotypes with cellular phenotypes. To overcome this limitation, we leveraged multi-modality single-cell sequencing, capturing the mutation with transcriptomes and methylomes in stem and progenitors from individuals with *DNMT3A* R882 mutated clonal hematopoiesis. *DNMT3A* mutations resulted in myeloid over lymphoid bias, and in expansion of immature myeloid progenitors primed toward megakaryocytic-erythroid fate. We observed dysregulated expression of lineage and leukemia stem cell markers. DNMT3A R882 led to preferential hypomethylation of polycomb repressive complex 2 targets and a specific sequence motif. Notably, the hypomethylation motif is enriched in binding motifs of key hematopoietic transcription factors, serving as a potential mechanistic link between *DNMT3A* R882 mutations and aberrant transcriptional phenotypes. Thus, single-cell multi-omics pave the road to defining the downstream consequences of mutations that drive human clonal mosaicism.

## INTRODUCTION

Somatic mutations have been recently identified ubiquitously across healthy tissues, indicating the presence of acquired clonal mosaicisms^1–6^. These mutations are pervasive across tissues such as the blood^7–17^, skin^5^, lung^2^ and esophagus^1,3^, and their prevalence increases with physiological aging. Importantly, somatic mutations in these clonal outgrowths overlap with recurrent drivers of cancer (for example, *DNMT3A*, *TP53, PIK3CA,* and *NOTCH1*)^1–5,8,18^, suggesting that cancer may arise from pre-malignant clonal outgrowths. Nevertheless, mutated cells are morphologically and phenotypically similar to their wildtype counterparts. This limits the ability to define the downstream transcriptional or phenotypic impact that may drive clonal outgrowth, and therefore prior studies in primary human tissue have largely focused on genetic characterization of clonal mosaicism.

Clonal mosaicism within the hematopoietic system serves as an informative model for this phenomenon, as recurrent drivers of myeloid malignancies (for example, *DNMT3A*, *TET2* and *ASXL1* mutations) have been detected in individuals without overt hematologic abnormalities^7–17^. This state, termed clonal hematopoiesis (CH), predisposes these individuals to an increased risk of developing myeloid malignancies, such as acute myeloid leukemias (AML) and myelodysplastic syndromes, and thus represents the earliest stages of neoplastic evolution^8,19–21^. Intriguingly, CH mutations also increase the risk of cardiovascular disease^11^ and progression of non-myeloid malignancies^11,22,23^, with early evidence supporting an aberrant immune microenvironment due to CH^8,24–26^. CH mutations have also been found in stem cell grafts, linked with idiopathic cytopenia in graft recipients^27^. CH mutations in certain genes (e.g. *DNMT3A*, *TP53*) endow a particularly strong fitness advantage in the context of stem cell transplantation, wherein the variant allele frequencies (VAF) markedly increase post-transplant compared to pre-transplant grafts^28,29^. These data suggest that certain CH mutations confer a particularly robust competitive advantage over non-neoplastic hematopoietic cells in stressed settings such as transplantation.

*DNMT3A*, which encodes a de novo DNA methyltransferase that catalyzes the methylation of cytosine bases in CpG dinucleotides, is by far the most frequently mutated gene in CH^7–10^. Consistently, *DNMT3A* mutations are considered an early event in AML^7^, and the hotspot variant at R882 constitute the majority of *DNMT3A* mutations in AML. The frequency of R882 variants is lower in CH, suggesting that these variants are particularly prone to progressing to AML through clonal evolution^12,30,31^. In vitro and murine models have suggested that *DNMT3A* R882 (or the murine R878 homologous residue) mutations result in a differentiation block and increased self-renewal in the hematopoietic stem cells (HSCs)^32–34^. Biochemically, *DNMT3A* R882 variants may exhibit a dominant negative effect^35,36^, resulting in the reduction of methyltransferase activity^36^. However, the study of *DNMT3A* mutations directly in human samples has been largely limited to MDS or AML, where confounding co-occurrence of other genetic alterations is common. Thus, CH presents a unique setting to interrogate the molecular consequences of *DNMT3A* mutations in non-malignant human hematopoiesis.

However, in CH as in other contexts of somatic mosaicism, mutated cells are admixed with wildtype cells^12,31^, limiting our ability to link genotype to phenotype using studies of bulk populations. Although recent fluidics methods for single-cell genotyping coupled with oligo-barcoded antibodies have begun to shed light on the phenotypic consequences of CH mutations^37^, these methods are limited to a small number of pre-defined cell surface markers. To overcome this limitation, we applied multi-omics single-cell sequencing to capture the mutational status of individual cells together with downstream epigenetic and transcriptional information^38,39^, thus enabling us to compare mutated cells with their wildtype counterparts from the same individuals, directly in primary human samples.

## RESULTS

### Genotyping of *DNMT3A* mutations in single-cell RNA-seq of CD34+ cells of human clonal hematopoiesis

As individuals with CH have normal blood production and thus meet no clinical criteria for assessments by bone marrow biopsy, progenitor-enriched samples with CH are scarce. However, we recently observed that CH is prevalent in patients with multiple myeloma (MM), and thus we interrogated a cohort of 136 MM patients with CH identified in hematopoietic progenitor cells collected for autologous stem cell transplant while in remission^40^. Given the known strong phenotypic impact of *DNMT3A* R882 mutations, we focused on four samples with these mutations and sufficiently high VAFs of >0.05 (range: 0.09-0.34) to enable profiling of large numbers of mutated cells with single-cell RNA-sequencing (scRNA-seq; see patient and sample data in **Extended Data Fig. 1a**; **Supplementary Table 1**). Notably, although CH mutations tend to have low VAFs, CH clones with higher VAFs have been frequently observed^8,10,41^. We further confirmed that no morphologic evidence of a myeloid neoplasm was seen in the bone marrow (**Supplementary Table 1**). Screening for additional mutations through a targeted myeloid panel^40^ showed only one additional mutation (patient CH03), consisting of a clonal (VAF = 0.5) heterozygous *TET2* nonsense mutation, which therefore likely arose first in the course of clonal evolution and serves as a background mutation for both the *DNMT3A* R882 mutated and wildtype cells.

We isolated viable CD34^+^ cells from these CH samples and performed Genotyping of Transcriptomes (GoT^38^), capturing scRNA-seq with targeted genotyping of the R882 codon (**Fig. 1a**). A total of 27,324 cells across CH samples were sequenced and included in the downstream analysis after quality filters (online methods, **Extended Data Fig. 1b**). Genotyping data were available for 6,430 cells of these 27,324 cells (23.5%) through GoT (**Extended Data Fig. 1a,c,d**). Notably, to overcome the challenge of accurate genotyping of the lowly expressed *DNMT3A* gene, we performed deeper sequencing and further optimized the original GoT analysis pipeline (IronThrone^38^, see online methods). This optimization included integrating unique molecule identifier (UMI) consensus assembly^42^, resulting in enhanced precision, with increased number of cells correctly assigned with only mutant or wildtype UMIs in a species mixing experiment (P < 10^−10^, Fisher exact test, **Extended Data Fig. 1e**). We also filtered the GoT UMIs based on their presence in the 10x gene expression library to determine the threshold for the number of supporting reads (online methods, **Extended Data Fig. 1f**). Mutated CD34^+^ cell frequencies ranged from 13% to 50%, comparable to the VAFs obtained through bulk sequencing of matched unsorted stem cell products (**Extended Data Fig. 1a,c**). Finally, to exclude additional genetic lesions, we performed copy number analysis with scRNA-seq data^43^ and identified no significant chromosomal gains or losses (**Extended Data Fig. 2a,b**).

**Figure 1.**
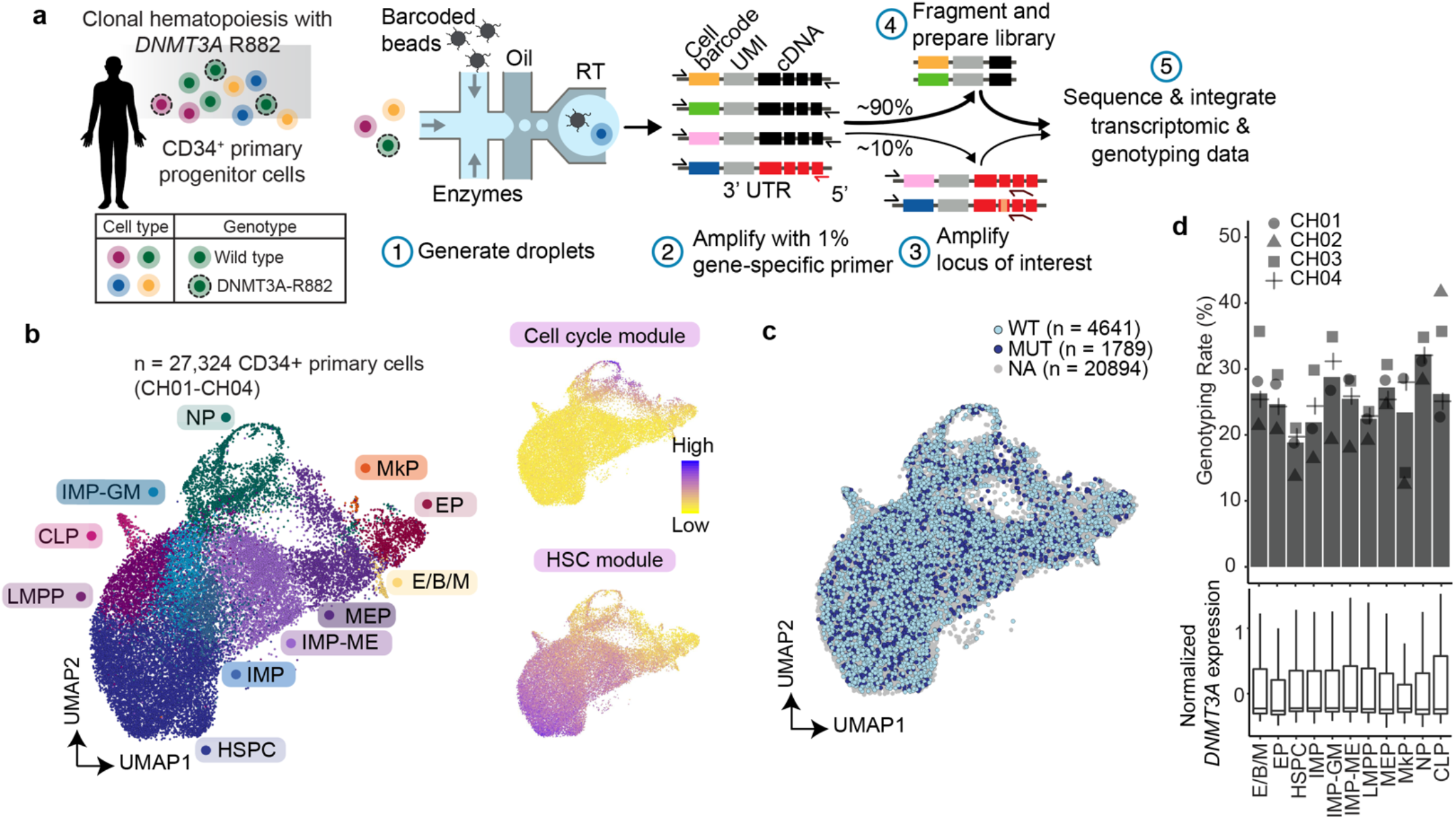
Genotyping of Transcriptomes demonstrates co-mingling of mutated and wildtype cells in *DNMT3A* R882-clonal hematopoietic differentiation. **a,** Schematic of GoT workflow. UMI, unique molecular identifier; UTR, untranslated region. **b,** Uniform manifold approximation and projection (UMAP) of CD34^+^ cells (n = 27,324 cells) from clonal hematopoiesis samples (n = 4 individuals), overlaid with cluster assignment (left); projections of cell cycle gene module (top right) or uncommitted hematopoietic stem cell (HSC) associated gene module score (bottom right, **Supplementary Table 2**). **c,** UMAP of CD34^+^ cells (n = 27,324 cells) with projected mutation status assignment for WT (n = 4,641 cells), *DNMT3A* R882 mutant (MUT; n = 1,789 cells) or unassigned (NA; n = 20,894 cells). **d**, Percent of genotyped cells per cluster for all samples (bars) and for each patient sample (points) (top) and normalized gene expression of *DNMT3A* per cluster (bottom). HSPC, hematopoietic stem progenitor cells; IMP, immature myeloid progenitors; IMP-ME, megakaryocytic-erythroid biased IMP; IMP-GM, granulo-monocytic biased IMP; LMPP, lympho-myeloid primed progenitors; CLP, common lymphoid progenitor; MEP, megakaryocytic-erythroid progenitors; E/B/M, eosinophil, basophil, and mast cell progenitors; EP, erythroid progenitor; MkP, megakaryocytic progenitor; NP, neutrophil progenitor; WT, wildtype; MUT, mutant; NA, not assignable.

To chart the differentiation of CD34^+^ progenitor cells in CH, we integrated data across the samples^44^ and clustered based on transcriptomic data alone, agnostic to the genotyping information (**Fig. 1b**, **Extended Data Fig. 3a**, online methods). Consistent with clinical data indicating normal hematopoietic production, we identified the expected progenitor subtypes, using previously annotated progenitor identity markers (**Fig. 1b**, **Extended Data Fig. 3b-d**, **Supplementary Table 2**)^45^. Furthermore, consistent with the fact that G-CSF mobilizes early stem and progenitor cells, we identified a large population of the earliest hematopoietic stem progenitor cells (HSPCs), as well as immature myeloid progenitor cells (IMPs), previously defined in a landmark scRNA-seq study^45^ as corresponding to the phenotypically-defined common myeloid progenitors (CMPs) and granulocyte-monocyte progenitors (GMPs). The high-throughput profiling by digital scRNA-seq enabled a higher resolution view of the IMPs, revealing a subcluster that exhibited markers of granulocyte-monocyte differentiation (IMP-GM) and a subcluster that exhibited markers of megakaryocytic-erythroid differentiation (IMP-ME, **Extended Data Fig. 4a,b**). Having established the progenitor identities, we then projected the genotyping information onto the differentiation map (**Fig. 1c, Extended Data Fig. 4c)**. No novel cell identities were formed by the *DNMT3A* mutations, consistent with the fact that patients with CH exhibit no overt peripheral blood count or morphologic abnormalities, Instead, we observed that mutated and wildtype cells co-mingled throughout (**Fig. 1c, Extended Data Fig. 4c**), highlighting the need for single-cell multi-omics to link genotypes with cellular phenotypes in CH. Importantly, the genotyping efficiency was balanced across the progenitor subsets, mitigating potential technical biases (**Fig. 1d, top**), consistent with no significant difference in *DNMT3A* gene expression within the CD34^+^ cell subsets (**Fig. 1d, bottom**).

### DNMT3A-mutated cells show lineage biases at key differentiation junctures

As previous data in murine and in vitro models have suggested that *DNMT3A* mutations may lead to a differentiation block^46,47^, we performed a differentiation pseudo-temporal (pseudotime) ordering analysis of the GoT data^48–50^. We found no significant global difference between wildtype and mutated cells (P = 0.70, linear mixed model, **Extended Data Fig. 4d** including per sample analysis, online methods), indicating that *DNMT3A* R882 mutations do not result in a significant global differentiation block in pre-cancerous human hematopoietic development. This finding is nonetheless consistent with findings in murine models, where even in the setting of homozygous *Dnmt3a* deletion, mutated cells do not exhibit self-renewal advantage in primary transplant experiments^47^, indicating that features of self-renewal advantage may not be overtly obvious in steady-state hematopoiesis. Although we did not observe a global differentiation block, we hypothesized that the *DNMT3A* mutated cell frequencies may vary across certain progenitor identities. For example, as *DNMT3A* R882 mutations are more frequently associated with myeloid rather than lymphoid neoplasms, we tested whether mutated cells may demonstrate a lineage bias toward myeloid versus lymphoid differentiation by examining lympho-myeloid primed progenitors (LMPP) and common lymphoid progenitors (CLP). Consistent with frequency biases seen in murine models for *DNMT3A* mutations^51^, mutated cells were enriched in myeloid biased cells versus early lymphoid progenitors (P < 0.001, linear mixed model, **Fig. 2a**). Moreover, these data are also consistent with previous results obtained with bulk, sorted populations from a *DNMT3A* I780T CH sample, which showed a lower VAF in mutated cell frequency in mature lymphoid cells (e.g. NK cells, B cells), compared to those in myeloid progenitor and mature cells^52^.

**Figure 2.**
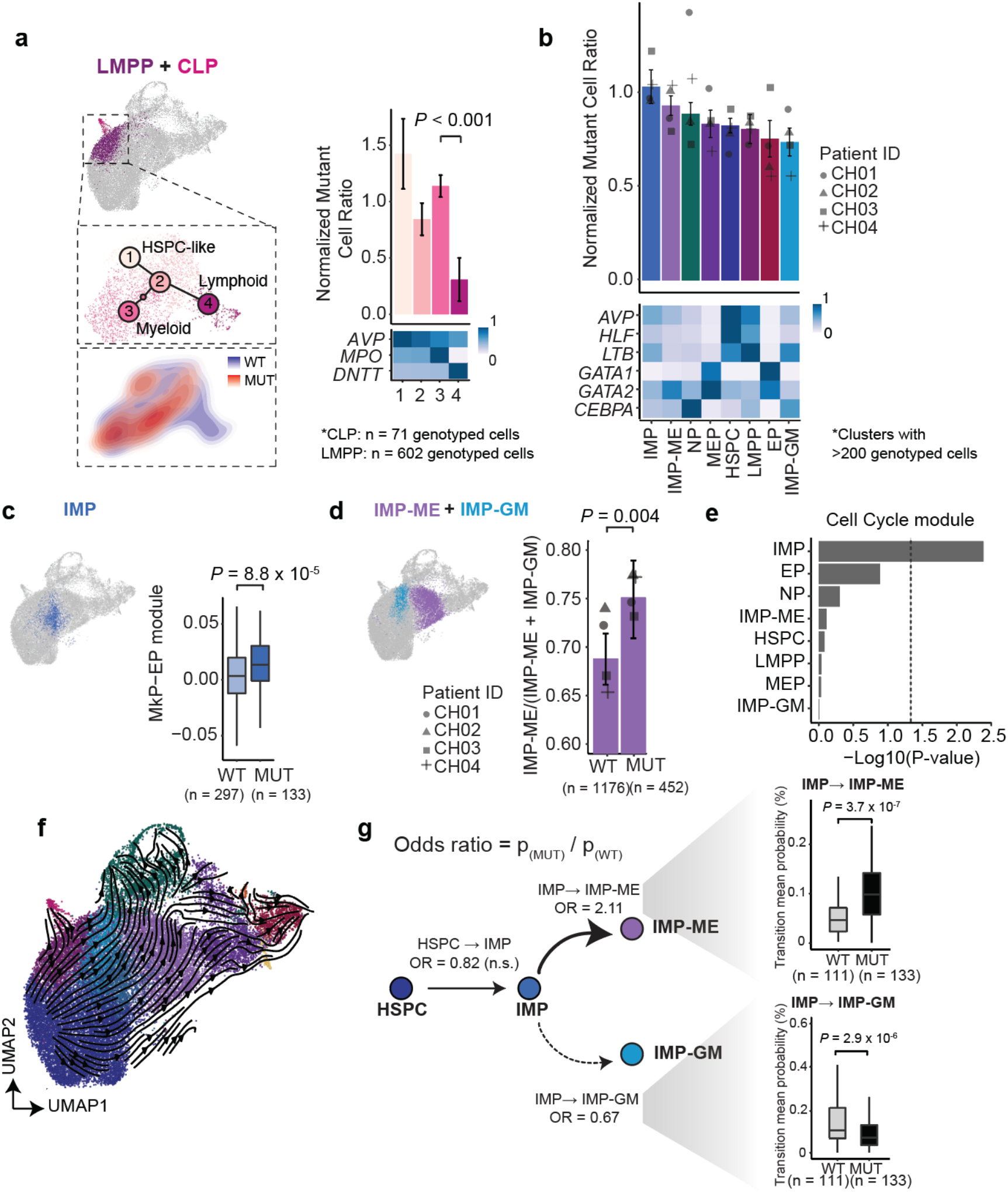
*DNMT3A* R882 mutated CH cells demonstrate distinct differentiation biases at key junctures. **a,** UMAP highlighting multi-lineage lympho-myeloid primed progenitors (LMPPs) and common lymphoid progenitors (CLPs); UMAP showing analytically isolated and re-clustered LMPPs and CLPs, showing branch point of divergence into myeloid versus lymphoid primed progenitors (left middle); UMAP showing the cell density of *DNMT3A* R882 MUT and WT cells (left bottom). The normalized frequency of mutant cells in subclusters for aggregate analysis of samples CH01-CH04 with mean ± s.d. of 100 downsampling iterations to 1 genotyping UMI per cell (right, downsampling performed to control for potential greater ability to detect the mutant heterozygous allele in cells with higher *DNMT3A* expression, see online methods). The heatmap at the bottom depicts representative lineage-specific genes for individual clusters. P-value was calculated from likelihood ratio test of LMM with/without cluster identity. **b,** Normalized frequency of *DNMT3A* R882 mutant cells in progenitor subsets with at least 200 genotyped cells. Bars show aggregate analysis of samples CH01-CH04 with mean ± s.d. of 100 downsampling iterations to 1 genotyping UMI per cell. Points represent mean of n = 100 downsampling iterations for each sample. Heatmap depicts representative lineage-specific genes for individual progenitor subsets. **c,** Megakaryocytic-erythroid module scores in wildtype versus mutant IMPs (**Supplementary Table 2**). P-value was calculated from likelihood ratio test of LMM with/without mutation status. **d,** Fraction of IMP-ME cells out of all biased IMP (IMP-ME + IMP-GM) cells in wildtype versus *DNMT3A* R882 mutant populations. P-value was calculated from proportions test. **e,** Cell cycle module scores in wildtype versus mutant progenitor subsets (**Supplementary Table 2**). P-values were calculated from likelihood ratio test of LMM with/without mutation status. **f,** RNA velocity field vectors overlaid on UMAP, demonstrating differentiation trajectories computed via scVelo (online methods). **g,** Schematic representation of the transition probabilities between HSPCs and IMP subsets from samples CH01-CH04 (right). Odds ratios (OR) were calculated as the ratio between *DNMT3A* R882 MUT and WT transition probabilities, as measured using RNA velocity. Single cell mean IMP → IMP-ME or IMP → IMP-GM transition probabilities between wildtype or *DNMT3A* R882 mutant cells, inset. P-values were calculated from likelihood ratio test of LMM with/without mutation status (see **Extended Data Fig. 6** for per-sample data).

To identify differentiation biases more broadly in *DNMT3A*-mutated CH, we evaluated the mutated cell frequencies across the different prevalent progenitor cell types (>200 genotyped cells). Of note, as cells may display variable expression of *DNMT3A* itself, we performed amplicon UMI down-sampling to exclude sampling biases given the heterozygosity of the mutated allele as a potential confounder for observed differences in mutated cell frequencies^38^. We observed that across samples, mutated cells were enriched in IMPs compared to the earliest HSPCs (P < 0.001, linear mixed model, **Fig. 2b**). Mutated IMPs also displayed an ME bias with an increase in the expression of an MkP-EP gene set^53^ (P = 8.8 × 10^−5^, linear mixed model, **Fig. 2c**, **Supplementary Table 2**, online methods), consistent with an increase in the proportion of IMP-ME to IMP-GM in mutant compared to wildtype cells (P = 0.004, proportions test, odds ratio of 1.38 (1.08 – 1.76), **Fig. 2d**). These data are in line with evidence of subtle erythroid abnormalities observed in CH via routine clinical assays (e.g. elevated red cell distribution width (RDW))^21^, and with our recent demonstration of increased HSC erythroid priming in a *Dnmt3a* knock-out murine model^54^.

Increased mutated cell frequency in a specific progenitor subtype can result from cell-type specific elevated proliferation^38^. We therefore first compared the expression of cell cycle genes^55^ between mutated and wildtype progenitors and found a modest increase in cell cycle gene expression only in mutated IMPs (P = 4.1 × 10^− 3^, linear mixed model, **Fig. 2e**, **Extended Data Fig. 5a**). Alternatively, increased mutated cell frequency in a given progenitor subtype, may stem from a change in transition rates into this cell state. To explore this hypothesis, we measured transition probabilities between progenitor subtypes with RNA velocity (online methods)^56,57^. The overall RNA velocity measurements demonstrated that these mobilized CD34^+^ cells follow the expected differentiation trajectories as described in normal human bone marrow hematopoiesis^53,58^ (**Fig. 2f**). Consistent with the hypothesis that transition rates contribute to the observed differentiation biases, we identified that the transition probability of mutated IMPs to become IMP-MEs was higher compared to that of wildtype cells (P = 3.7 × 10^−7^, linear mixed model, **Fig. 2g,** see **Extended Data Fig. 6a** for per sample comparison), whereas the transition probability of mutated IMPs to IMP-GMs was diminished (P = 2.9 × 10^− 6^, linear mixed model, **Extended Fig. 6b**). These analyses thus orthogonally confirmed ME-biased differentiation of *DNMT3A*-mutated CD34^+^ human progenitors, as was also revealed by the gene set expression analysis (**Fig. 1c**).

### Gene expression changes in *DNMT3A* mutated cells include leukemia stem cell genes, and are linked to proinflammatory signatures and putative dysregulated transcription factor activity

To identify the transcriptional dysregulation that may underlie the observed differentiation biases, we performed differential gene expression analysis between mutated and wildtype progenitors within each progenitor cell type. Differential expression (DE) analysis of mutated versus wildtype HSPCs revealed 88 dysregulated genes (**Fig. 3a,** 68-122 differentially expressed genes in each progenitor subset, see **Supplementary Table 3** for results for each progenitor subset; batch-aware permutation test where mutated and wildtype labels are permuted only within the same sample, see online methods). Of note, to ensure that the analysis was not dominated by a single sample, we down-sampled the number of mutated and wildtype cells from each sample to maintain equal representation in the progenitor subset DE analysis. To test the robustness of our approach further, we also determined DE by an alternative linear mixed model framework, in which we explicitly modeled samples as a random effect variable, and identified a high degree of concordance between the two statistical frameworks (**Extended Data Fig. 7a**, **Supplementary Table 3**, online methods).

**Figure 3.**
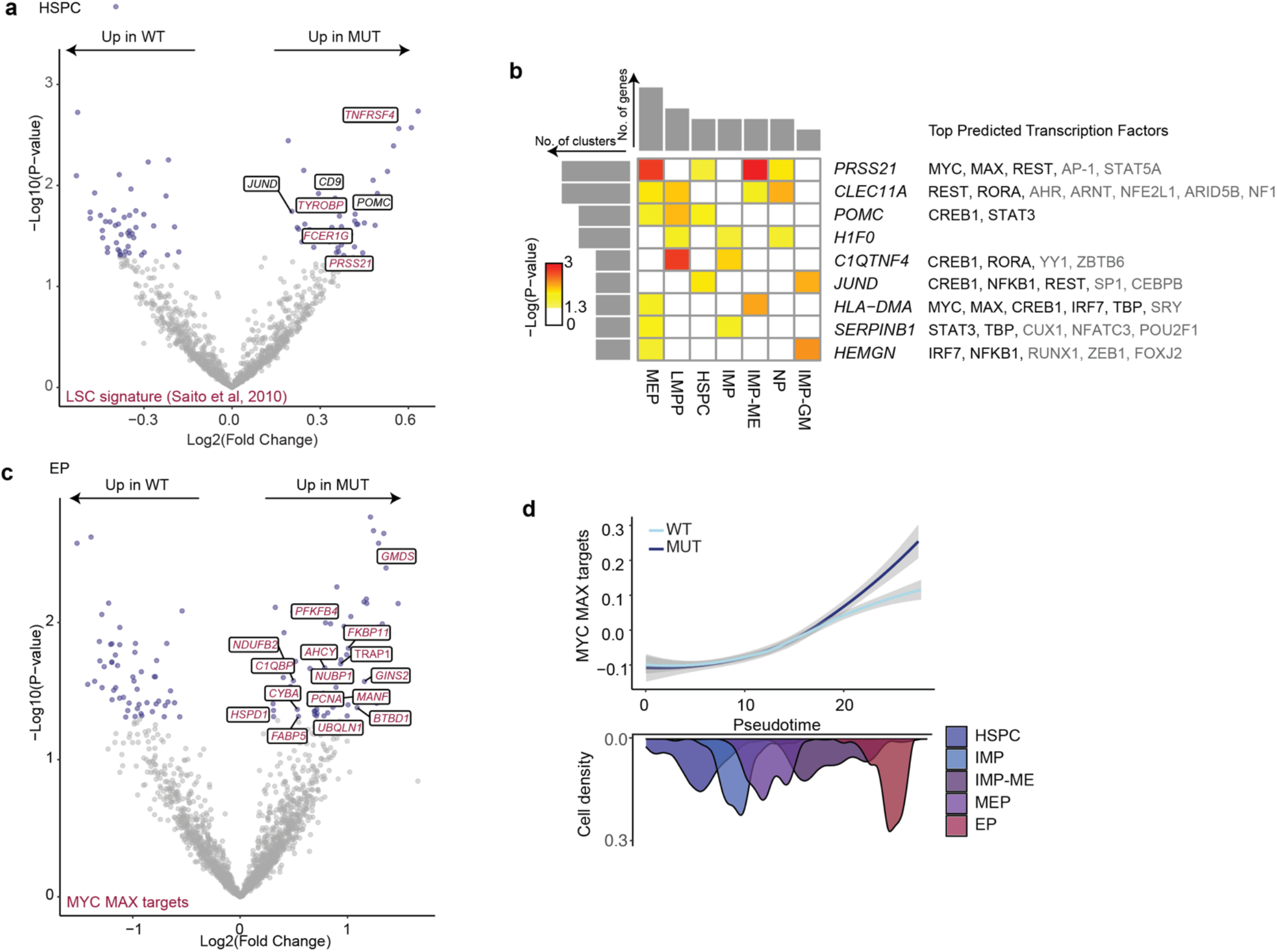
Differential gene expression analysis between mutated and wildtype cells reveals markers of lineage aberrancies and dysregulated MYC activity. **a,** Differentially expressed (DE) genes between *DNMT3A* R882 mutant and wildtype hematopoietic stem progenitor cells (HSPC) via permutation test (online methods). Genes highlighted in red represent DE genes overlapping with 58 genes upregulated on acute myeloid leukemia stem cells (LSC) compared to normal HSCs (P = 9.3 × 10^−5^). P-value was calculated by hypergeometric test. **b,** Heatmap of upregulated genes in *DNMT3A* mutant cells compared to wildtype cells, in at least two cell clusters (P < 0.05, permutation test). Histograms show numbers of upregulated genes in each cluster (top) and numbers of clusters per upregulated gene (left). Next to the genes are listed putative TFs (TRANSFAC) with black indicating the TFs that overlap for more than one recurrent DE gene. **c,** Differentially expressed genes between *DNMT3A* R882 mutant and wildtype EPs via permutation test. Pathway enrichment of MSigDB CGP gene sets shows enrichment of Benporath MYC MAX targets (FDR-adjusted P-value = 0.01) and Coller MYC targets (FDR-adjusted P-value = 0.01, see **Supplementary Table 4** for complete gene set enrichment results against the MSigDB CGP dataset). P-values were calculated from hypergeometric test with FDR (Benjamini-Hochberg) correction. **d,** Local regression of normalized expression levels as a function of pseudotime of MYC/MAX targets (differentially upregulated in **Fig. 3c**) for WT and *DNMT3A* R882 mutant (MUT) cells. Shading denotes 95% confidence interval. Histogram shows cell density of clusters included in the analysis, ordered by pseudotime.

DE genes included, for example, the upregulation of *CD9* in the early mutated HSPCs (**Fig. 3a**, **Supplementary Table 3**). CD9 expression is closely linked with megakaryocytic-priming^59,60^ and platelet activation^61–63^, thus providing further support for the ME bias of *DNMT3A* mutated progenitors. These data are also in line with a lower degree of thrombocytopenia observed in patients with *DNMT3A* mutated versus wildtype AML^64,65^ and thrombocytosis in a murine model of this mutation^66^. We further observed an enrichment of genes previously associated with leukemia stem cells (LSCs)^67^ in mutated HSPCs, including *PRSS21*, *FCER1G, TYROBP,* and *TNFRSF4*, mapping these dysregulated genes to the nascent neoplastic process (P = 9.3 × 10^−5^, hypergeometric test, **Fig. 3a**, **Supplementary Table 3**). *FCER1G, TYROBP* and *TNFRSF4*, are known to be involved in proinflammatory signaling^68–76^, consistent with previous reports suggesting that CH clones display enhanced proinflammatory signatures^24,26,41,77–81^. In another example, we identified upregulation of the pro-survival oncogene *PIM2*, downstream of STAT signaling, in mutated LMPPs, recently implicated as a target for eradicating chemotherapy-resistant chronic myeloid leukemia stem cells^82^ (**Supplementary Table 3**).

Nine genes were upregulated in more than one progenitor subset (**Fig. 3b**, **Supplementary Table 3**). This analysis highlighted mediators of cell-to-cell interactions, such as a regulator of the inflammatory network *C1QTNF4*^83,84^. We also identified *CLEC11A* (also known as stem cell growth factor (SCGF)), which has been implicated as a hematopoietic growth factor^85,86^, including in the setting of hematopoietic stress such as irradiation and transplantation^85,87^. This finding is consistent with published murine data showing a 6.75-fold increase of *Clec11a* in transplanted *Dnmt3a* KO cells compared to wildtype cells^88^. Thus, overexpression of *CLEC11A* by *DNMT3A*-mutated progenitors may provide a potential mechanism for marked expansion of CH clones upon transplantation^28,29,89–93^. Genes upregulated in more than one progenitor subset were associated with putative transcription factors^94^, identifying recurring TFs (highlighted in black, **Fig. 3b**), including MYC and its cofactor MAX, as well as the inflammatory NFKB and STAT transcription factors and interferon regulatory factor IRF7, consistent with proinflammatory networks in CH clones^24,26,77,80,81^.

To more broadly identify dysregulated pathways, we performed a gene set enrichment analysis of the differentially upregulated genes (**Fig. 3c**, **Supplementary Table 4**)^95,96^. The top significantly enriched pathways (FDR < 0.2) included MYC targets in the mutated erythroid progenitors (FDR-adjusted P = 0.01, **Fig. 3c**). Notably, we observed the enrichment of two independent MYC target gene sets, including a MYC signature that was downregulated with monocytic differentiation in an HSPC differentiation cell line model^97,98^. Consistently, MYC has been demonstrated to be a critical factor specifically for erythropoiesis^99–101^, and may thus contribute to the observed ME bias (**Fig. 2c,d,g**). Of interest, *DNMT3A* mutation driven MYC target expression increased during differentiation along the erythroid lineage (**Fig. 3d**), despite no increase in *MYC* gene expression itself in the mutated progenitors (**Extended Data Fig. 7b**), suggesting that its transcriptional output as a transcription factor is differentially increased in mutated cells. Other enriched pathways included targets of cell cycle regulator E2F in LMPPs (FDR-adjusted P = 0.057, **Supplementary Table 4**). Altogether, these findings suggest a focused dysregulation in TF activity that may orchestrate the observed lineage and transcriptional perturbations in the premalignant stages of hematopoietic neoplasia.

### Single-cell multi-omics integrating somatic genotyping, methylome, and transcriptome profiling reveals patterns of *DNMT3A* mutation hypomethylation

To directly decipher the underlying link between mutated *DNMT3A*-induced DNA hypomethylation and the observed altered transcriptional regulatory networks in CH, we profiled CD34^+^ progenitors from the same individuals (from samples CH02 and CH04 where additional material was available) with multi-modality single-cell sequencing capturing DNA methylation (DNAme)^102^, scRNA-seq (Smart-seq2^103^), and targeted *DNMT3A* genotyping^39^ (n = 528 cells after quality filtering, **Fig. 4a,b**, **Extended Data Fig. 8a-c**, online methods). As expected, these scRNA-seq data identified the major progenitor identities as those demonstrated by the 10x platform, albeit at a lower resolution given fewer cells (**Fig. 4b,** left, **Extended Data Fig. 8b**). Of these 528 cells, genotyping data were available for 441 cells (**Fig. 4b**, right, 84% cells genotyped). This multi-modal profiling uniquely enabled us to compare the methylation status of mutated and wildtype cells from the same individuals, showing a decrease in DNAme in CpG islands even in this relatively heterogeneous CD34^+^ population (CGI, P = 5.72 × 10^−3^, linear mixed model, **Fig. 4c**), consistent with the finding that *DNMT3A* mutated AMLs have lower methylation of CGI compared to *DNMT3A* wildtype AMLs^104^. While enhancers have been demonstrated to be particularly impacted by *DNMT3A* loss in the setting of AML^105^, these relatively CpG-poor regions have lower coverage in standard enzymatic methyl-seq (EM-seq)^106^ or reduced representation bisulfite sequencing (RRBS) with a single restriction enzyme Msp1. We therefore increased the capture of enhancer regions through double restriction-enzyme Msp1 and HaeIII digestion^107^ and identified marked hypomethylation of enhancer regions^108^ (P = 7.29 × 10^−8^, linear mixed model, **Fig. 4d**) as well as global hypomethylation in *DNMT3A* R882 cells compared to wildtype cells (P = 2.92 × 10^−3^, linear mixed model, **Extended Data Fig. 8c-d**, online methods). Thus, we demonstrated that the methylation of regulatory regions is affected by *DNMT3A* R882 mutations in human CH. Interestingly, prior in vitro studies suggested that CpH sites may be hypermethylated in *DNMT3A* R882H. Our data revealed no significant difference, and an opposite trend (**Extended Data Fig. 8e),** further highlighting the significance of examining primary human cells.

**Figure 4.**
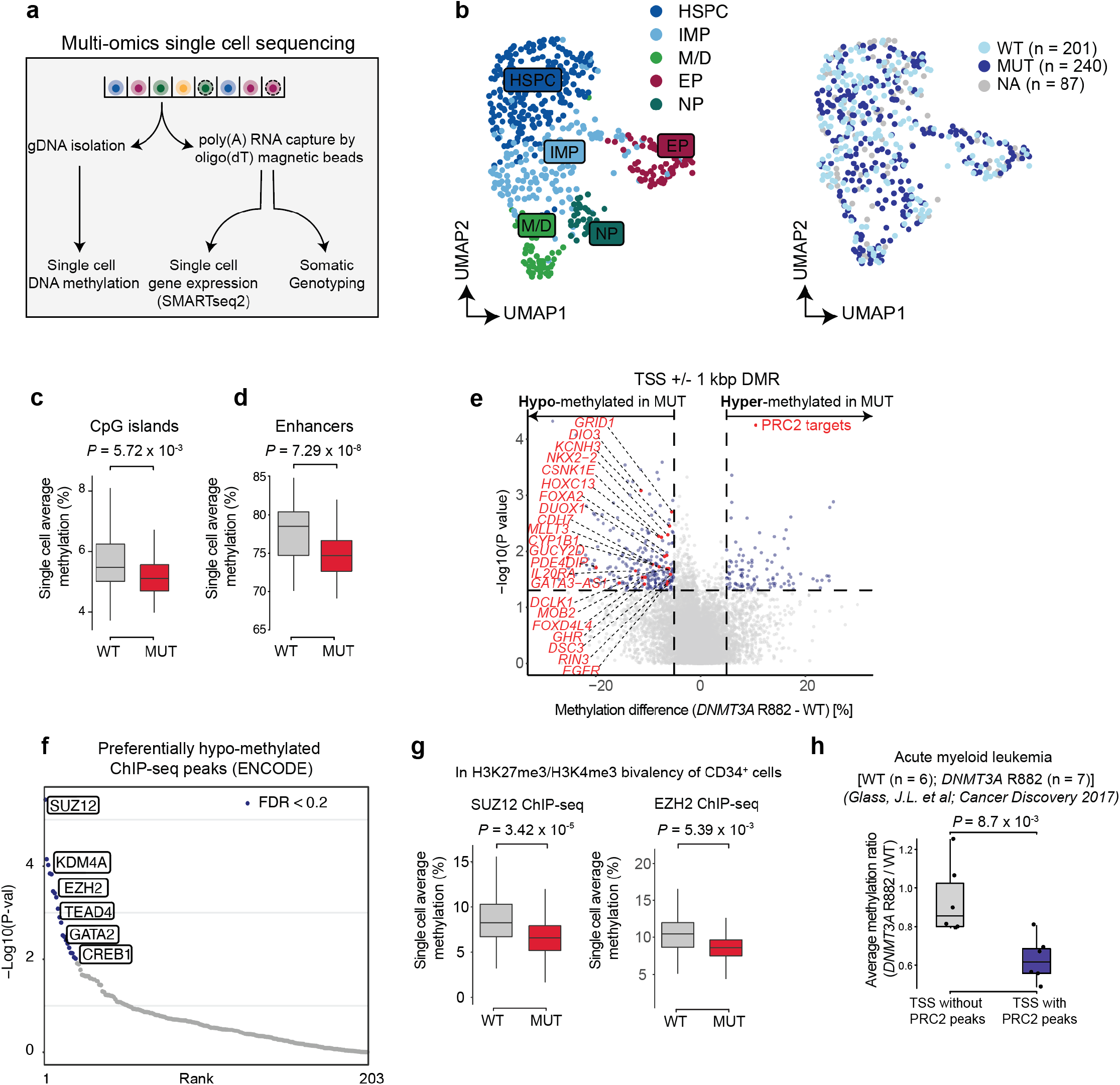
*DNMT3A* R882 promotes selective hypomethylation of PRC2 targets in human hematopoiesis. **a,** Schematic representation of the single-cell multi-omics platform that captures methylome, transcriptome, and somatic genotype status**. b,** UMAP dimensionality reduction (n = 528 cells) showing the assigned progenitor identities (left) or the assigned genotype (right) from available samples CH02 and CH04. (**c-d**) Average single cell methylation at CpG islands **c,** and enhancers **d,** from double digest experiments (online methods). P-values from likelihood ratio test of LMM with/without mutation status. **e,** Differentially methylated promoters between wildtype and *DNMT3A* R882 mutant hematopoietic progenitors. P-values from generalized linear model (GLM) to account for global hypomethylation in *DNMT3A* mutated cells and identify regions of preferential hypomethylation (online methods). Red dots indicate significantly hypomethylated Benporath PRC2 and EED target genes (MSigDB C2: CGP gene sets). **f,** Differentially hypomethylated ChIP-seq peaks (ENCODE hg38 Tf clusters) ranked by P-value. P-values from a GLM to account for global hypomethylation in *DNMT3A* mutated cells and identify regions of preferential hypomethylation. **g,** Single cell average methylation at ChIP-seq peaks (ENCODE hg38 Tf clusters intersected with bivalent peaks (H3K27me3, H3K4me3) from human CD34^+^ hematopoietic progenitor cells) for either SUZ12 (left) or EZH2 (right). P-values from likelihood ratio test of LMM with/without mutation status. **h,** Comparison of AML samples with/without *DNMT3A* R882 showing *DNMT3A* mutant-to-wildtype ratio of methylation at TSS overlapping PRC2 ChIP-seq peaks or non-overlapping CpG rich TSS as control. P-value from two-sided Wilcoxon rank sum test. HSPC, hematopoietic stem progenitor cells; IMP, immature myeloid progenitor; NP, neutrophil progenitor; M/D, monocytic/dendritic cell progenitors; EP, erythroid progenitor; WT, wildtype; MUT, mutant; NA, not assignable.

Differentially methylated regions (DMR) analysis identified 269 promoters to be significantly hypomethylated considering the observed global hypomethylation (P < 0.05 and at least 5% loss in methylation, **Fig. 4e**, **Extended Data Fig. 8f**, **Supplementary Table 5,** see online methods for statistical modeling to identify promoters with preferential hypomethylation that explicitly models samples as a variable). Gene set enrichment analysis of these genes identified enrichment of targets of the PRC2 (FDR-adjusted P < 0.2, GSEA with MSigDB C2: CGP gene set, **Fig. 4e**, **Supplementary Table 6**, online methods). As an orthogonal approach, we performed differential methylation analysis of chromatin immunoprecipitation sequencing (ChIP-seq) peaks (ENCODE database^109^) that overlap with TSS regions. This approach also identified the targets of PRC2 components SUZ12 and EZH2 to be differentially hypomethylated (**Fig. 4f**), as well as that of GATA2, involved in ME differentiation. As ENCODE ChIP-seq tracks reflect aggregation across several cell types, we validated that preferential hypomethylation specifically impacted regions marked by H3K27me3, H3K4me3 bivalency in human hematopoietic progenitors, by intersecting the ENCODE ChIP-seq tracks with bivalent peaks in CD34^+^ cells^110^ (**Fig. 4g**, **Supplementary Table 7**, for per-sample data see **Extended Data Fig. 8g**). This finding is consistent with previous data showing that germline gain-of-function mutations in *DNMT3A* result in the reciprocal *hyper*methylation of PRC2 targets, leading to premature differentiation programs^111^. Furthermore, PRC2 targets exhibit significant overlap with previously reported methylation canyons, shown to undergo preferential hypomethylation upon *Dnmt3a* loss^112^ (98% of canyons harbored a PRC2 target compared with 16% of canyons harboring peaks of a size-matched set of random genomic intervals, P < 10^−10^, Fisher exact test)^113^. Notably, while gene expression changes in PRC2 targets were not observed between mutated and wildtype cells from the GoT data (P = 0.42, linear mixed model, **Extended Data Fig. 8h**), this may be expected given that PRC2-repressed genes that gain DNA methylation may only switch between different silencing states. Nonetheless, DNA methylation of PRC2 targets has been shown to reinforce gene silencing^114–116^, and thus mutated *DNMT3A* mediated hypomethylation of PRC2 targets may poise mutated progenitors to aberrant reactivation of stem cell maintainers, as seen in a PRC2 deficient mouse model^117^.

Finally, to determine whether CH hypomethylation of PRC2 targets persists through progression to AML, we compared the methylation status of PRC2 targets (online methods) between *DNMT3A* R882 mutated AML (n = 7) and *DNMT3A* wildtype AML (n = 6, both groups with *NPM1* mutations^105^, **Supplementary Table 8**). We found that compared with *DNMT3A* wildtype AML, *DNMT3A* R882 mutated AML demonstrated preferential hypomethylation at promoters of PRC2 targets compared to promoters with similar CpG content (P = 0.0087, online methods, **Fig. 4h**, **Extended Data Fig. 8i**). To determine whether the preferential hypomethylation of PRC2 targets may be robust against various co-occurring mutations, we compared the methylation rates of PRC2 targets in *DNMT3A* wildtype (n = 122) versus *DNMT3A* R882 mutated AML (n = 9) with heterogeneous mutation status from The Cancer Genome Atlas (TCGA)^118^ and identified similar results as observed in the *NPM1*-mutated AML (**Extended Data Fig. 8j**). These results demonstrated that mutated *DNMT3A*-mediated hypomethylation of PRC2 targets is maintained through evolution to AML, further supporting it as a potential mechanism for enhanced self-renewal, from clonal hematopoiesis to frank malignancy.

### *DNMT3A* R882 displays differential methyltransferase activity as a function of CpG flanking sequence

We hypothesized that mutated *DNMT3A* R882 may further display differential methyltransferase activity, depending on the flanking sequence context of the CpG dinucleotide^119,120^. Indeed, CpGs within DMRs defined CpG motifs that are particularly hypomethylated (disfavored) in mutated versus wildtype human CD34^+^ cells (online methods, **Fig. 5a**, **Extended Data Fig. 9a**). Of note, CpGpT was particularly associated with hypomethylation (**Fig. 5a**, **Extended Data Fig. 9a**), consistent with in vitro enzymatic studies of DNMT3A R882 variants^119,120^ (**Extended Data Fig. 9b,c**). Importantly, this CpG flanking motif was enriched in the binding motifs of specific TFs expressed in hematopoietic progenitors (**Fig. 5b**). These included key regulators of hematopoiesis such as MYC/MAX, whose activities are known to be negatively impacted by DNA methylation of their binding motifs^121,122^, and were found to have increased target expression in mutated cells (**Fig. 3c,d**). Other key transcription regulators included HIF1A (and its cofactor ARNT), whose binding is facilitated by demethylation of the binding motif^123^; HIF1A/ARNT are critical factors for HSC quiescence, through maintenance of the anaerobic glycolysis-dependent metabolic activity in the bone marrow niche^124–130^. USF1/2 were also among the highlighted TFs, which have been shown to regulate chromatin architecture in erythroid differentiation and the beta-globin locus^131,132^. In further support for a model in which preferential hypomethylation of the specific sequence motif underlies transcriptional dysregulation, we observed enrichment of the hypomethylated CpG flanking sequence in regions surrounding genes upregulated in mutated HSPCs and EPs (**Fig. 5c**, **Extended Data Fig. 9d-f**).

**Figure 5.**
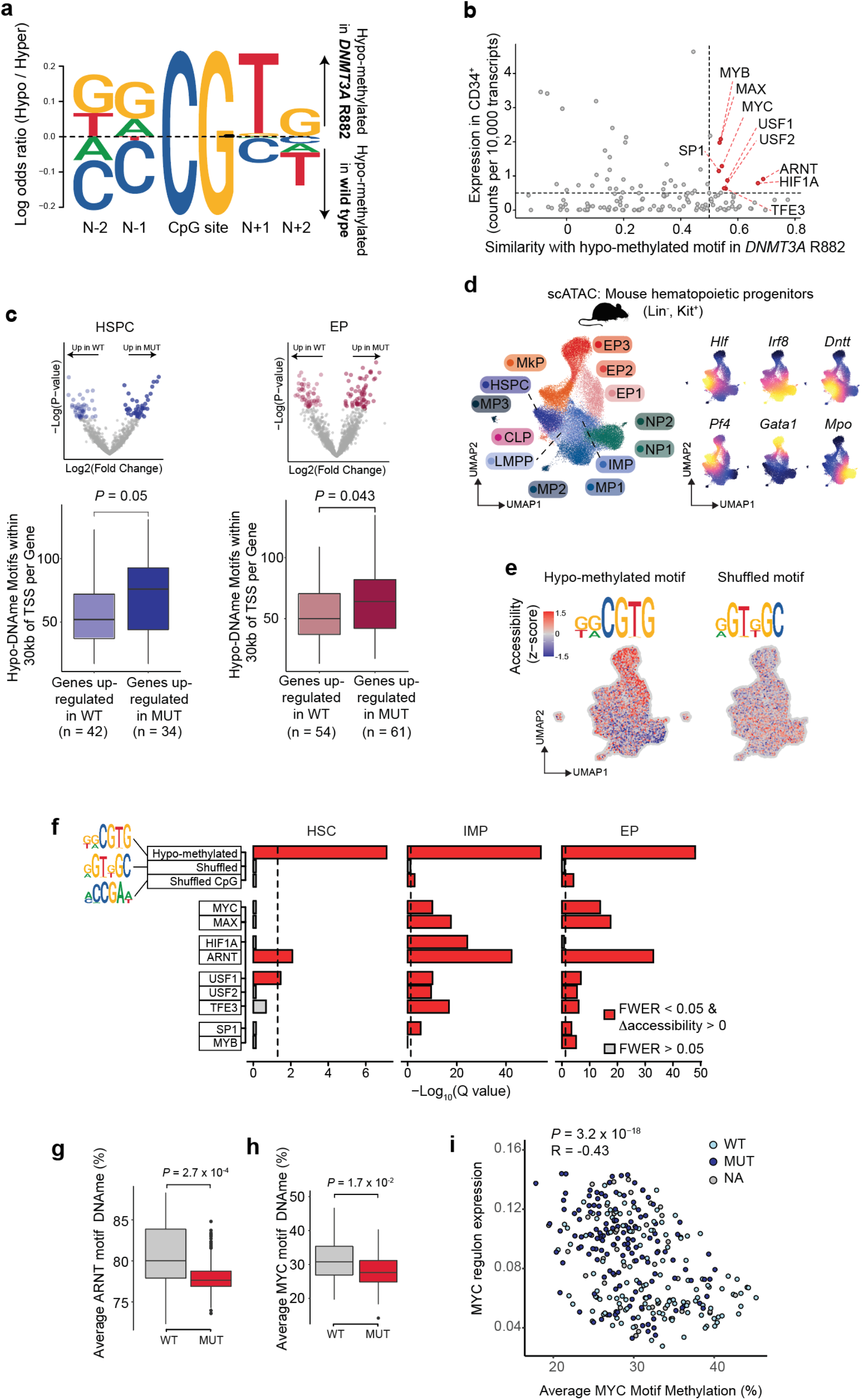
*DNMT3A* R882 displays flanking sequence specificity associated with MYC binding motif. **a,** Motif logo for the odds ratio of base frequency of the flanking positions (N-1, N-2, N+1, N+2) of CpG sites. Odds ratios were calculated based on the flanking regions of CpG sites hypomethylated or hypermethylated in *DNMT3A* R882 mutant compared with wildtype hematopoietic progenitors (online methods)**. b,** Similarity score between the hypomethylated motif of *DNMT3A* R882 (**Fig. 5a**) and TF binding motifs in the HOCOMOCO v11 collection of human TF binding motifs. Relevant transcription factors with expression level in HSPCs > 0.5 and motif similarity > 0.5 are labeled. **c,** Frequencies of *DNMT3A* R882 hypomethylated motif within 30 kb of TSS of the differentially expressed genes between MUT and WT cells in HSPCs and EPs (identified in GoT data, **Fig. 3a,c,** see **Extended Data Fig. 9d** for other progenitor subsets, **Extended Data Fig. 9e** for 10 kb and 50 kb of TSS, and **Extended Data Fig. 9f** for data accounting for CpG content). P-values were calculated by Wilcoxon rank sum test. **d,** UMAP dimensionality reduction of murine wildtype (n = 3 mice) and *Dnmt3a* R878H (n = 3 mice) Lin^−^, Kit^+^ snATAC-seq data showing progenitor cluster annotation and representative progenitor gene marker accessibility (n = 46,496 cells). **e,** UMAP showing accessibility deviation as calculated with chromVar for hypomethylated motif (left) and shuffled motif (right, z-scores). **f,** Bonferroni FWER-adjusted P-values for accessibility changes between wildtype and *Dnmt3a* R878H cells by progenitor identities for hypomethylated motif and negative control shuffled motifs (with/without CpG), as well as binding motifs of the TFs identified in **Fig. 5b**. **g,** Comparison of single cell average methylation of ARNT binding motifs (intersected with ARNT ChIP-seq peaks, ENCODE hg38 Tf clusters) between wildtype and *DNMT3A* R882 mutant hematopoietic progenitor cells. P-values from likelihood ratio test of LMM with/without mutation status. **h,** Comparison of single cell average methylation of MYC binding motifs (intersected with MYC ChIP-seq peaks, ENCODE hg38 Tf clusters) between wildtype and *DNMT3A* R882 mutant hematopoietic progenitor cells. P-values from likelihood ratio test of LMM with/without mutation status. **i,** Relative expression per cell (AUC) of MYC downstream targets inferred using the SCENIC package (online methods) as a function of average MYC motif methylation. Correlation coefficient R calculated using Pearson’s Correlation. P-value derived from GLM. HSPC, hematopoietic stem progenitor cells; MP, multipotent progenitors; IMP, immature myeloid progenitors; LMPP, lympho-myeloid primed progenitors; CLP, common lymphoid progenitor; EP, erythroid progenitor; MkP, megakaryocytic progenitor; NP, neutrophil progenitor.

To validate the impact of mutated *DNMT3A* on TF activation, we collected Lin-, c-Kit+ hematopoietic stem and progenitor cells from mice with and without *Dnmt3a* R878H (the murine R882H equivalent; no. of mice = 3 in each cohort)^51^. While recent progress has been made in single-cell chromatin binding assays^133–135^, the ability to determine the weaker signal of TF binding in single cells remains a challenge. We therefore performed a chromatin accessibility assay, shown to be a reliable surrogate for determining TF activity^136^, on single nuclei (n = 46,496 cells, **Fig. 5d**, **Extended Data Fig. 10a-d**). Confirming our findings in human CH, we found that the accessibility of the *DNMT3A* R882-specific hypomethylated motif was increased in R878H cells, across clusters, including in HSPCs, and particularly in EPs (**Fig. 5e,f**, **Extended Data Fig. 10e-g**), whereas shuffled versions of the hypomethylated motif, with or without a CpG, displayed lower difference in accessibility between mutated and wildtype progenitors. Candidate TFs with high similarities scores in their binding motif with the hypomethylated motif, including MYC/MAX, HIF1A/ARNT, USF1/2, displayed enhanced accessibility in R878H compared with wildtype progenitors, across multiple progenitor subsets (**Fig. 5f**, **Extended Data Fig. 10g**). The myeloid progenitors were particularly impacted, whereas the lymphoid progenitors showed little to no significant difference in accessibility for these TF binding motifs (**Extended Data Fig. 10g)**, suggesting overactivity of these TFs may play a role in the myeloid differentiation bias. While *Dnmt3a* R878H HSPCs displayed a more modest increase in chromatin accessibility, this may be due to the global open chromatin in stem cells reducing the ability to measure specific enrichments^137,138^. Overall, as chromatin accessibility has been demonstrated to accurately reflect TF activity^136^, these data provided further evidence for the model in which the *DNMT3A* mutation enhances the activity of TFs whose binding motifs are prone to hypomethylation through enrichment in the hypomethylated sequence motif. This model then provides the basis of enhanced MYC/MAX target gene expression in the *DNMT3A* mutated cells observed in the GoT data (**Fig. 3c,d**), despite no expression increase in the *MYC* gene itself (**Extended Data Fig. 7b**). With respect to PRC2 targets, although hypomethylation of PRC2 target genes were observed, we observed no differential increase in expression in mutated cells (**Extended Data Fig. 8h**) and no enhanced accessibility of PRC2 targets in the mutated cells from mouse snATAC-seq data (**Extended Data Fig. 10h**).

As further confirmation of our proposed model, we found that HIF1A/ARNT and MYC/MAX binding motifs were hypomethylated in CH mutated cells compared to wildtype progenitors in the single-cell multi-omics data (P = 2.7 x 10^−4^ and P = 1.7 × 10^−2^, respectively, linear mixed model, **Fig. 5g,h**). Moreover, as MYC targets were upregulated in CH mutated cells in the GoT data, we leveraged our single-cell multi-omics approach to directly link the expression of MYC/MAX targets with the level of DNA methylation of MYC/MAX target promoters within the same cells (see online methods). Indeed, the expression of MYC/MAX target genes was negatively correlated with mean methylation of their binding sites (P = 3.2 × 10^−18^, generalized linear model, **Fig. 5i**), consistent with prior studies indicating that hypomethylation of binding motifs enhances MYC binding^121,122,139,140^. Thus, our single-cell multi-omics profiling provides a potential model for the observed transcriptional aberration in human *DNMT3A* mutated CH, supporting enhanced fitness of *DNMT3A* mutated cells via selective hypomethylation of key hematopoietic TF binding motifs.

### *DNMT3A*-mutated CH bone marrow sample corroborates results from stem cell graft CH samples

To confirm that the findings we observed in the CH samples were generalizable to CH not exposed to G-CSF or prior chemotherapy, we obtained a bone marrow sample from a patient without any underlying hematologic disorders with a *DNMT3A* R882H mutation (CH05). We sorted for CD34^+^ cells and performed GoT as we had done for CH01-CH04 samples (n = 5,770 cells). Although a low genotyping efficiency limited the comparisons between mutated and wildtype cells within the same sample (n = 687 genotyped cells), this sample consisted of mostly mutated cells with a high VAF (0.4), enabling a direct comparison to previously published healthy control CD34^+^ bone marrow cells (n = 39,082 cells, **Supplementary Table 9**, online methods)^141,142^. We batch-corrected and integrated across the samples as previously described^44^ (**Fig. 6a,b, Extended Data Fig. 11a-e**). We first determined whether the bone marrow CH IMPs may display the lineage biases as previously observed in the CH01-CH04 samples. Consistent with those results, the IMPs from CH05 demonstrated skewing toward the ME versus GM state, compared to the control bone marrow CD34^+^ cells (**Fig. 6c, Extended Data Fig. 12a-c**). Next, we assessed the progenitor-specific differentially expressed genes identified in the CH01-CH04 samples and confirmed the expected increased or decreased expression for the differentially upregulated or downregulated genes in mutated cells, respectively, in CH05 progenitors, compared to control progenitors (data for HSPCs and EPs in **Fig. 6d, Extended Data Fig. 12d,e**, see other progenitors in **Extended Data Fig. 12f**). Furthermore, we observed an enrichment of the MYC/MAX target genes in the CH05 progenitors compared to the control progenitor cells (**Fig. 6e**), again most pronounced within the EPs. Intriguingly, the CH05 cells integrated evenly across progenitor subsets with control CD34^+^ cells except for a subcluster of EPs (EP2, **Fig. 6a-b, Extended Data Fig. 12g**). We suspected that the MYC/MAX target gene expression may be particularly impacted in this aberrant cluster and identified this to be the case (**Fig. 6e, right**). While the low genotyping efficiency limited our ability to make within cluster mutated versus wildtype comparisons in this sample, we were able to confirm across clusters the increased expression of differentially upregulated genes identified in more than one progenitor subset (**Extended Data Fig. 12h-j**, genes from **Fig. 3b**). Lastly, to test whether CD9 protein expression was impacted by the upregulation of the gene expression observed in the mutated HSPCs from CH01-CH04, we incorporated protein expression in this sample through CITE-seq^143^. As CD9 expression has been linked with megakaryocytic differentiation priming^59,60^, we examined CD9 protein expression in the in the early CD34^+^, CD38^low^ hematopoietic stem and progenitor cells along the megakaryocytic differentiation trajectory and observed an increased CD9 expression in mutated compared with wildtype cells (**Extended Data Fig. 12k,l**).

**Figure 6.**
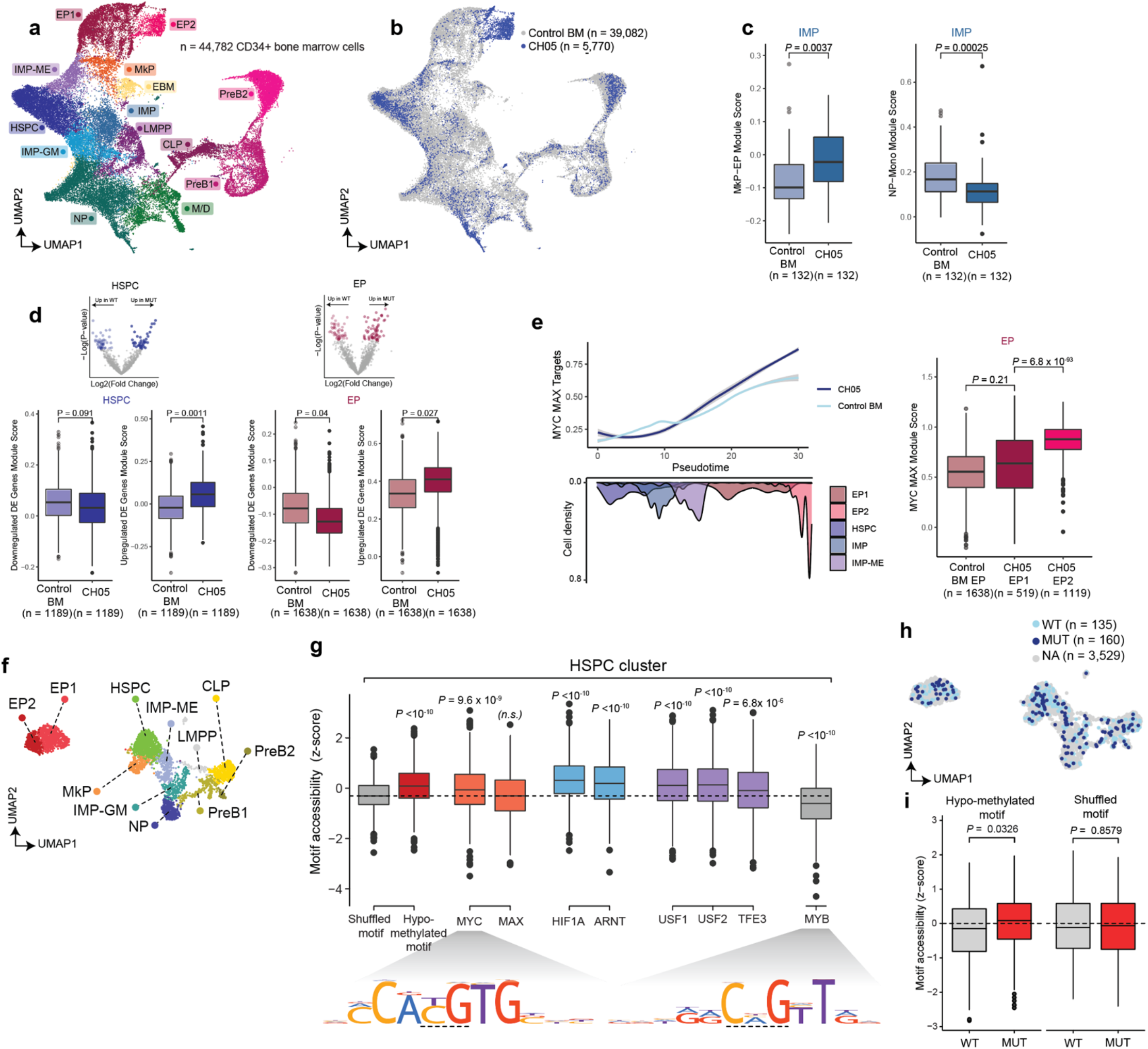
Bone marrow clonal hematopoiesis progenitor cells display megakaryocytic-erythroid differentiation bias, MYC target gene expression, and enhanced accessibility for the R882 hypomethylated motif. **a,** UMAP of CD34^+^ cells (n = 44,782 cells) for scRNA-seq data from a clonal hematopoiesis sample (CH05) and previously published five control bone marrow samples (BM01-05), labeled with cluster assignments. **b,** UMAP of CD34^+^ cells (n = 44,782 cells) labeled with CH (n = 5,770) or control (n = 39,082) status. **c,** Megakaryocytic-erythroid module scores in control versus CH IMPs (left, **Supplementary Table 2**) granulocytic-monocytic module scores in control versus CH IMPs (right, **Supplementary Table 2**). P-values were calculated from likelihood ratio test of LMM with/without CH status. **d,** Module scores for differentially down- or up-regulated genes in mutant *DNMT3A* HSPCs and EPs (identified in GoT data, **Fig. 3a,c**) in control versus CH HSPCs and EPs. **e,** Local regression of normalized expression levels as a function of pseudotime of MYC/MAX targets (differentially upregulated in **Fig. 3c**) for control and *DNMT3A* R882 CH cells. Shading denotes 95% confidence interval. Histogram shows cell density of clusters included in the analysis, ordered by pseudotime. Boxplot shows comparison of module scores between control and CH cells within the two EP clusters. P-value calculated from likelihood ratio test of LMM with/without CH status. **f**, UMAP dimensionality reduction of CD34^+^ cells (n = 3,824 cells) for snATAC-seq data from a clonal hematopoiesis sample (CH05) depicting the cell cluster assignment and cell type labels. **g,** Motif accessibility z-scores for shuffled, hypo-methylated motif and relevant transcription factors for the HSPC cluster (n = 788 cells). P-values correspond to Wilcoxon rank sum test between accessibility of the shuffled motif and the indicated motif. **h,** UMAP projection of genotype assignment for WT (n = 135 cells) and MUT (n = 160 cells). **i,** Motif accessibility z-score comparison for either hypo-methylated or shuffled motifs between WT (n = 135 cells), MUT (n = 160 cells). P-values were calculated by Wilcoxon rank sum test. HSPC, Hematopoietic stem and progenitor cell; IMP-ME, immature myeloid progenitor with megakaryocytic/erythroid bias; IMP-GM, immature myeloid progenitor with granulocyte/monocyte bias; LMPP, Lymphoid-myeloid pluripotent progenitor; MkP, Megakaryocyte progenitor; NP, Neutrophil progenitor; CLP, Common lymphoid progenitor; Pre-B1/2, Pre-B cell progenitor; EP1/2, Erythroid progenitor.

To test the chromatin accessibility of TF motifs (as a surrogate for TF activity) that bear high similarity to the *DNMT3A* R882 hypomethylated motif directly in this CH sample, we extended GoT to the 10x Multiome (ATAC+RNA) platform and applied it to sorted CD34^+^ nuclei (**Fig. 6f**, **Extended Data Fig. 13a-c**, n = 3,824 nuclei, note that the transcriptome data failed QC metrics and was not used downstream). As genotyping efficiency depends on mRNA abundance, the lower mRNA abundance in nuclei limited genotyping. We therefore again took advantage of the high VAF (~80% cells are mutant) and showed that across these cells, the accessibility of the hypomethylated motif – as well as those of MYC/MAX, HIF1A/ARNT, USF1/2/TFE3 – was increased compared to a shuffled motif and that of MYB, which may serve as an additional negative control (**Fig. 6g**). The accessibility of the hypomethylated motif increased with erythroid differentiation but not with lymphoid differentiation, consistent with the importance of these highlighted TFs in erythroid differentiation (**Extended Data Fig. 13d,e**). Finally, even within the limited number of genotyped cells, we observed that the accessibility of the hypomethylated motif was increased overall in the mutated cells compared to the wildtype-enriched population (**Fig. 6h,i**). In summary, these findings in a bone marrow *DNMT3A*-mutated CH sample, not complicated by exposure to G-CSF or prior chemotherapy, corroborated the findings in samples CH01-CH04, suggesting that the comparisons between mutated and wildtype cells within the same individuals are indeed robust to the potentially confounding extrinsic factors and are largely generalizable to steady-state *DNMT3A* R882-mutated CH.

## Discussion

We present an unbiased profiling of the downstream effects of somatic driver mutations in clonal mosaicism of normal human tissue, focusing on *DNMT3A* mutations in clonal hematopoiesis. Hitherto, extensive genetic profiling across normal tissues has been performed to document the striking mosaicism that result from pervasive age-related acquisition of somatic mutations^1–5^. For example, a landmark study of morphologically normal skin from the eyelids of four individuals identified ~140 mutations per square centimeter^5^. Importantly, while these studies have demonstrated that mutations in cancer drivers are particularly prevalent^5^, the downstream effects of cancer driver mutations that enable clonal outgrowths in normal human tissue are largely unknown.

Similarly, CH is a prevalent phenomenon in physiological hematopoietic aging fueled by driver mutations linked with myeloid neoplasms. However, the downstream consequences of these mutations in normal human hematopoietic progenitors are largely unknown. Previous studies leveraged rare germline mutations in small cohorts of patients to study the downstream perturbations of these mutations^104,111^. For example, by examining mature blood cells from an individual with Tatten-Brown-Rahman Syndrome (TBRS) due to a germline *DNMT3A* R882H mutation, with a sibling control^104^, the Ley group demonstrated focal hypomethylation, including of CpG islands, consistent with our findings. More recently, the Goodell group studied the effects of *DNMT3A* R771Q mutation by transforming primary cells into a lymphoblastoid cell line (LCL) from an early embryonal mosaic individual^144^, demonstrating significant overlap in hypomethylated regions in these *DNMT3A* R771Q LCLs and *DNMT3A* mutated AML.

Nonetheless, we previously lacked the ability to directly compare mutated and wildtype progenitors in human CH in their native context. Specifically, two obstacles challenge the study of CH mutation impact directly in primary patient samples. First, CH specimens with enriched human hematopoietic progenitors are scarce, as individuals with CH have no current clinical indication for a bone marrow biopsy. To circumvent this limitation, we pursued an alternative approach to profile CH mutated cells in stem cell graft products obtained from a cohort of MM patients in remission^145^ and identified one *DNMT3A* R882H CH bone marrow specimen without G-CSF exposure or a potentially confounding cancer diagnosis to validate our findings. Second, mutated cells are admixed with wildtype in the hematopoietic progenitor pool and are morphologically and phenotypically indistinct. Thus, mutated cells cannot be isolated from wildtype cells for downstream analysis. We overcame this challenge by leveraging single-cell multi-omics that enabled us to profile the transcriptomes and epigenomes, together with the genotype information, of these single cells.

The application of the GoT approach^38^ enabled high-resolution mapping of *DNMT3A* R882 mutated cells to the hematopoietic differentiation tree to reveal differentiation skewing, even before clinically observable changes in blood counts. We observed a myeloid over lymphoid bias, consistent with prior murine studies^51^, and the strong association of this genotype with myeloid versus lymphoid neoplasms. We further identified expansion of mutated IMPs and ME-biased IMPs. Enrichment of mutated cells in IMPs was linked with a specific increase in proliferation compared to wildtype cells. Notably, myeloid-bias has been linked with proinflammatory signaling^64,146^, and thus a proinflammatory state in mutated HSPCs (i.e. as evidenced by the overexpression of *TNFRSF4*, *TYROBP*, *FCER1G*) may also contribute to the enrichment of mutated cells in IMPs. Mutated IMPs further displayed a megakaryocytic-erythroid lineage bias, with enhanced transition probability of mutated IMPs to differentiate into IMP-MEs, consistent with our previous study in *Dnmt3a* KO mouse model^54^, as well as a *Dnmt3a* R878H model showing increased platelet counts^64^.

As *DNMT3A* R882-induced changes in DNAme are globally distributed across the genome, we sought to understand how stochastic DNAme changes can be translated into deterministic outputs, especially with respect to differentiation skews. We found that the DNMT3A R882 variants displayed a CpG sequence motif specificity, disfavoring CpGs with T at the N+1 position, consistent with deep enzymology assays^119^. Notably, this hypomethylated CpG flanking motif bore high similarity to the binding motifs of key hematopoietic TFs, such as MYC/MAX, HIF1A/ARNT, USF1/2, providing a mechanistic model for enhanced MYC activity observed in our GoT data. This model was supported by mouse *Dnmt3a* R878H and, critically, human CH bone marrow data in which snATAC-seq of hematopoietic progenitors revealed enhanced accessibility of the hypomethylated motif and importantly of the MYC/MAX, HIF1A/ARNT, USF1/2 binding motifs. The accessibility changes associated with the hypomethylated motif were specifically pronounced in myeloid versus lymphoid progenitors, suggesting that these molecular consequences may play a role in differentiation biases. Furthermore, our single-cell multi-omics platform further enabled us to identify that cells with hypomethylation of MYC/MAX binding motifs showed increased expression of their transcriptional targets within the same cells, consistent with previous reports that demonstrated the negative impact on MYC activity imparted by the methylation of its binding motif^121,122,139,140^. These data revealed how modest, global, stochastically distributed DNAme changes can be translated into phenotypic skews. Through differences in the enrichment of CpG flanking sequence density of TF DNA binding motifs, subtle global DNAme changes affecting hundreds of binding sites can modulate TF output to result in reshaping of the differentiation landscape^54^.

We further identified preferential hypomethylation of PRC2 targets. While the relationship between PRC2-mediated histone methylation and DNA methylation is not fully understood, DNA methylation may serve to “lock in” gene silencing with a mechanism with more robust mitotic inheritance^147^. PRC2 targets in stem cells include pluripotency/stemness genes^148–150^, and are enriched for bivalent H3K27me3/H3K4me3 marks^151,152^, suggesting that PRC2 results in “poising” rather than in complete silencing at those sites. In contrast, more differentiated cells reinforce gene silencing by increasing the length of H3K27me3 domains, or through complementary silencing mechanisms including DNA methylation^114–116^. Thus, while PRC2 targets are broadly suppressed in stem cells, some leaky transcription may still occur, compared to PRC2 targets that have also underwent DNA methylation. This nuanced model posits that PRC2 targets DNA hypomethylation in *DNMT3A* mutated progenitors, may allow for their re-activation in response to stimuli, as another candidate mechanism for enhanced self-renewal through de-repression of stem cell programs. As activation of stem cell markers such as those repressed by the polycomb group proteins have been implicated in endowing cancer with stem-like properties^153^, our data points to poising of PRC2 targets as a potential mechanism for enhanced stem cell renewal upon malignant transformation. While PRC2 deficiency has been reported to lead to overexpression of stem cell maintainers such as *HoxC4* and inhibitors of differentiation such as *Sox7* and *Id2* in a murine model (*Eed* KO), as well as relative expansion of LT-HSC^117^, *Eed* KO cells also showed reduced competitive repopulating capacities with pro-apoptotic predisposition^117^. These data suggest that PRC2 target activation of self-renewal requires cooperation of an oncogenic TF such as MYC to counterbalance the proapoptotic effects and support clonal expansion in *DNMT3A* R882 cells. In support of this model, a recent work in mice demonstrated that while *Ezh2* KO itself had little impact on hematopoiesis (likely due to redundant homologs), *Ezh2* KO together with a compounding oncogenic driver (*Nras* G12D) promoted myeloid malignancy with activation of stemness genes^154^. Interestingly, *Nras* G12D alone promoted GM over ME bias, but in the double *Ezh2* KO, *Nras* G12D mutated model, hematopoiesis was shifted toward ME over GM, suggesting that the PRC2 aberrations may indeed play a role in the observed ME bias (in addition to the better-established role of MYC in ME differentiation)^154^.

A potential limitation of our study of stem cell grafts is the exposure to G-CSF used in stem cell mobilization from patients with MM (of note, patients were not subject to other mobilization agents, such as CXCR4 antagonists or cyclophosphamide). Nonetheless, our analyses uniquely compared mutated and wildtype cells within the same sample, which were equally subjected to G-CSF. Indeed, our CH05 bone marrow aspirate sample from an individual with CH and no cancer diagnosis confirmed the major findings of the study, showing that comparing mutated versus wildtype cells from the same individuals is robust to the potential extrinsic confounders. For example, although G-CSF stimulates granulocytic differentiation and proliferation^155^, we were still able to capture the megakaryocytic-erythroid bias in the early mutated progenitors. Importantly, G-CSF is especially effective in mobilizing quiescent murine HSCs, without inducing proliferation^156^.

Interestingly, in the context of cell line models of *DNMT3A* R882, G-CSF induced a differentiation block in vitro in one study^34^ and GM-CSF masked the proliferative effects of the mutation in another^157^. Although these results were observed in cell lines, and thus the applicability to human CH is less clear, these data suggest that G-CSF may serve as a confounder. In this context, our validation of the major findings in a CH sample without exposure to G-CSF is of particular importance.

Another limitation results from the incomplete capture of the heterozygous allele in our GoT cDNA amplicon method due to low expression (median of 1 amplicon per genotyped cell, range 1-4 UMIs per cell). This is likely to result in misclassification of some mutated cells as wildtype cells. Nonetheless, as this is expected to diminish mutation-specific signals, the mutation-specific aberrations reported herein may likely have an even stronger effect size. Another limitation of the study is the sample size, due to the rarity of available samples. In this context, it is important to note that intensive profiling of a small number of samples (e.g. mutational profiling of normal eyelid samples from four individuals^5^ or epigenetic profiling of one TBRS patient with germline *DNMT3A* R882H mutation^104^) have shown that fundamental insights can be gained from these cases, directly in human samples. Our single-cell multi-omics profiling of thousands of progenitors, directly comparing mutated and wildtype cells within the same individuals, thus enabled us to highlight reproducible gene expression perturbations and epigenetic underpinnings, that were supported by evidence from published reports and murine data.

Altogether, we report the first direct examination of the molecular consequences of *DNMT3A* R882 mutations in primary CD34^+^ cells in human CH. These studies allowed us to directly superimpose the differentiation topographies of mutated and wildtype hematopoietic progenitors, co-existing within the same individuals. We identified key epigenetic and transcriptional aberrations that reshape the differentiation topography and contribute to clonal expansion in the most nascent stage of neoplasia. These data also demonstrate the power of emerging single-cell multi-omics methods^158–161^ to pave the road towards defining how mutations drive normal tissue mosaicism in human somatic evolution.

## Acknowledgments

The work was enabled by the Weill Cornell Epigenomics Core and Flow Cytometry Core. We thank Dr. Ari Melnick (Weill Cornell Medicine) for a critical review of the manuscript. A.S.N. is supported by the Burroughs Wellcome Fund Career Award for Medical Scientists and National Institutes of Health Director’s Early Independence Award (DP5 OD029619-01). N.D. is supported by a F30 Predoctoral Fellowship from the NHLBI of the National Institutes of Health (F30HL156496). N.D. and R.M.M. are supported by a Medical Scientist Training Program grant from the National Institute of General Medical Sciences of the National Institutes of Health under award number T32GM007739 to the Weill Cornell/Rockefeller/Sloan Kettering Tri-Institutional MD-PhD Program. F.I. is supported by the American Society of Hematology Fellow-to-Faculty Scholar Award. R.C. is supported by Lymphoma Research Foundation and Marie Skłodowska-Curie fellowships. D.A.L. is supported by the Burroughs Wellcome Fund Career Award for Medical Scientists, Valle Scholar Award, and the National Institutes of Health Director’s New Innovator Award (DP2-CA239065). This work was also supported by the National Heart Lung and Blood Institute (R01HL145283) and the National Human Genome Research Institute, Center of Excellence in Genomic Science (RM1HG011014).

## METHODS

### Patient samples

The study was approved by the local ethics committee and by the Institutional Review Board (IRB) of Weill Cornell Medicine, University of Chicago and Dana Farber Cancer Institute conducted in accordance to the Declaration of Helsinki protocol. All patients provided informed consent. Cryopreserved G-CSF mobilized stem cell grafts (without additional mobilizing agents such as plerixafor or cyclophosphamide) from patients in remission for multiple myeloma, with documented *DNMT3A* R882 mutations were retrieved after interrogating a cohort of 136 patients with CH^40^. See **Supplementary Table 1** for clinical information.

Cryopreserved grafts were thawed and stained using standard procedures (10 min, 4°C) with the surface antibody CD34-PE-Vio770 (clone AC136, lot# 5180718070, dilution 1:50, Miltenyi Biotec) and DAPI (Sigma-Aldrich). Cells were then sorted for DAPI-negative, CD34^+^ cells using BD Influx at the Weill Cornell Medicine flow cytometry core.

### Mouse Models

All animals were housed at Memorial Sloan Kettering Cancer Center (MSKCC). All animal procedures were completed in accordance with the Guidelines for the Care and Use of Laboratory Animals and were approved by the Institutional Animal Care and Use Committees at MSKCC. The *Dnmt3a* R878H mouse model has been described previously^51^, and was crossed to the *Tal1*-creERT2 transgenic model to allow for inducible control of the R878H mutation within the hematopoietic system^162^. To induce recombination of the conditional alleles, age and gender-matched 10-16 week old *Tal1*-creERT2 control mice and *Dnmt3a* R878H *Tal1*-creERT2 mice were treated with tamoxifen (4 mg/kg/day; Cayman Chemical, Ann Arbor, Michigan) for 2 doses, separated 2 days apart. The mice were sacrificed 4-8 weeks after tamoxifen-induction. Primary mouse bone marrow (BM) cells were isolated into cold phosphate-buffered saline (PBS), without Ca^2+^ and Mg^2+^, and supplemented with 2% bovine serum albumin (BSA) to generate single cell suspensions. Red blood cells (RBCs) were removed using ammonium chloride-potassium bicarbonate (ACK) lysis buffer, resuspended in PBS/2% BSA, and filtered through a 40μm cell strainer. Total nucleated cells were quantified by Vi-Cell XR cell counter (Beckman Coulter, Brea, CA) and used for downstream data production.

### Genotyping of Transcriptomes (GoT)

Genotyping of Transcriptomes was performed as previously described^38^. The standard 10x Genomics Chromium 3’ (v.3 chemistry) libraries were carried out according to manufacturer’s recommendations for the generation of scRNA-seq libraries (**Fig. 1a**). At the cDNA amplification step, 1 μL of 1 μM spike-in primer (5’ – GAGGTCAAACTCCATAAAGCAGGGC– 3’) was added to increase the yield of *DNMT3A* cDNA. After cDNA amplification and cleanup with SPRI beads, 25% of the cDNA underwent the standard 10x protocol per manufacturer recommendations. The unused cDNA was stored and 10% was subsequently used for targeted genotyping. For locus-specific amplification (GoT), two serial PCRs were performed with nested reverse primers (5’ –CTTATGGTGCACTGAAATGGAAAGGG – 3’ and 5’ – CCTTGGCACCCGAGAATTCCAGGTTTCCCAGTCCACTATACTGACG – 3’) and the generic forward SI-PCR were used to amplify the site of interest from the cDNA template (10 PCR cycles each). The second locus-specific reverse primer contains a partial Illumina TruSeq Small RNA read 2 handle and a locus-specific region to allow specific priming. The SI-PCR oligo (10x Genomics) anneals to the partial Illumina TruSeq read 1 sequence, preserving the cell barcode (CB) and unique molecule identifier (UMI). After these rounds of amplification and SPRI purification to remove unincorporated primers, a third PCR was performed with a generic forward PCR primer (P5_generic, 5’ – AATGATACGGCGACCACCGAGATCTACAC – 3’) to retain the CB and UMI together with an RPI-x primer (Illumina) to complete the P7 end of the library and add a sample index (6 cycles). The targeted amplicon library was subsequently spiked into the remainder of the 10x library to be sequenced together on a NovaSeq (Illumina). The cycle settings were as follows: 28 cycles for read 1, 98 cycles for read 2, 8 cycles for i7 and 8 cycles for i5 sample index.

### 10x scRNA-seq data processing, alignment, cell-type classification and clustering

10x data were processed using Cell Ranger (v3.0.1) with default parameters. Reads were aligned to the human reference sequence hg19. The genomic region of interest for genotyping was examined to determine how many UMIs with the targeted sequence were present in the conventional 10x data. The Seurat package (v.3.1) was used to perform unbiased clustering of the CD34^+^ sorted cells from patient samples^163^. In brief, for individual datasets, cells with UMI < 200 or UMI > 3 median absolute deviations from the median UMI, or mitochondrial gene percentage > 20%, were filtered. The data were log-normalized using a scale factor of 10,000. Before clustering, the individual datasets (CH01-CH04) were integrated and underwent batch-correction within Seurat, which implements canonical correlation analysis and the principles of mutual nearest neighbor^44^. Recommended settings were used for the integration (30 canonical correlation vectors for canonical correlation analysis in the FindIntegrationAnchors function and 30 principal components for the anchor weighting procedure in IntegrateData function). Following integration, potential confounders (specifically, number of UMIs per cell, proportion of mitochondrial genes, and patient sex) were regressed out of the data before principal component analysis was performed using variable genes using recommended settings (i.e. top 2000 variable genes using variance stabilizing transformation)^44^. The first statistically significant 30 principal components were used as inputs to the UMAP algorithm for cluster visualization^164^. Clusters were manually assigned on the basis of differentially expressed genes using the FindAllMarkers function using default settings (using all genes that are detected in a minimum of 25% of cells in either of the two comparison sets as input, and log-transformed fold change of 0.25 as the threshold). We identified 20 clusters in the integrated data, which were annotated according to canonical lineage markers identified previously in single-cell RNA-seq data of normal hematopoietic progenitor cells^53^. These clusters were collapsed into 11 main progenitor subsets based on expression of levels of these canonical markers (**Extended Data Fig. 3b,c**). Pseudotime analysis was performed using the Monocle3 R package using recommended parameters (v.0.2.1, **Extended Data Fig. 4d**) ^50^. In order to specify the initial cluster of the pseudotime trajectory, we identified the cluster with the highest expression level of the HSPC gene module (**Fig. 1b**, **Supplementary Table 2**). The Slingshot R package (v.1.6.1) was used to isolate the minimum spanning tree for the LMPP and CLP subset of cells (**Fig. 2a**) with default parameters.

### IronThrone GoT for processing targeted amplicon sequences and mutation calling

Analysis of the GoT library was carried out as described previously^38^. Briefly, amplicon reads were assessed for presence of the primer sequence and the expected sequence between the primer and the mutation site. Reads were also assessed for matching to the cell barcode list of the 10x dataset. A mismatch of 20% was allowed for all sequence matching steps. Only UMIs with at least 2 or more supporting reads were retained for final genotyping assignments. A few key improvements to our IronThrone pipeline (v.2.1) are detailed below.

First, parallelization was implemented to increase runtime efficiency for larger sequencing libraries^165^. The amplicon library of paired reads was shuffled and subsetted into smaller groups of reads (default 125,000 reads/group). Then, the original IronThrone algorithm was run on each one of these groups. This step has been parallelized using both GNU Parallel tools for local interactive operation, as well as options for Slurm-managed high-performance compute clusters. Output tables from these runs are finally concatenated by cell barcode.

Second, we improved the UMI counting of the amplicon reads by removing ‘pseudo’-UMIs introduced by PCR and sequencing errors (that would result in a false increase in the number of UMIs). Based on previously published work^42^, we implemented a network-based UMI collapsing algorithm to aggregate amplicon reads that likely originated from the same UMI in the original 10x library. Briefly, pairwise Levenshtein distances were calculated between all UMIs paired within a single cell barcode, and “matches” between UMIs were identified as UMI pairs with a Levenshtein distance below a predetermined threshold (default = ceiling(0.1 * UMI length), or 2 bases for a 12 base UMI). The UMI with the greatest number of matched UMIs was determined to be the initial UMI. The number of supporting reads for these UMI groups was summed together and attributed to that initial UMI with the most matches. This process was then repeated for the UMI with the next highest number of matches until no additional collapsing was possible. This improved pipeline was applied to the previously-described species mixing experiment^38^, demonstrating a significant improvement in the removal of aberrant genotyping UMIs (see Results, **Extended Data Fig. 1e**).

Following UMI collapse, genotype assignment of individual UMIs was conducted as described previously with majority rule of supporting reads for wildtype or mutant status. Rare UMIs with an equal number of mutant and wildtype reads were removed as ambiguous. Additionally, to remove reads that result from PCR recombination^38^, UMIs in the amplicon library that match UMIs of non-*DNMT3A* genes in the gene expression library were discarded. Of note, the latter likely PCR-recombination events were associated with lower number of read per UMI compared with UMIs in the amplicon library that matched *DNMT3A* UMI in the gene expression library (**Extended Data Fig. 1f**). We leveraged this observation, and retained UMIs without a corresponding associated gene in the gene expression library, so long as their read count was above the 80^th^ percentile of read counts for non-*DNMT3A* genes. Finally, single cells were assigned mutant or wildtype genotype status as follows: cells with one or more mutant UMIs were assigned as mutant cells, and cells with 0 mutant UMIs and at least one wildtype UMI were assigned as wildtype. While the genotyping information is derived from transcribed molecules and may be affected by the capture of transcripts from wildtype versus mutant alleles of heterozygous mutations, the frequency of mutant cells as determined by GoT using all cells that harbor at least one UMI yielded values that were similar to that determined by bulk DNA exon sequencing (**Extended Data Fig. 1c**).

### Mutant cell frequency analysis

To exclude the possibility that variable *DNMT3A* expression may impact the ability to detect mutant alleles and thereby impact mutated cell frequency in distinct progenitor subsets, we down-sampled all cells to a single amplicon UMI prior to mutation calling for calculating mutant cell frequencies (**Fig. 2a,b**). An equal number of cells from each sample CH01-CH04 (n = 83 cells for LMPP + CLP (**Fig. 2a**) and n = 978 cells for analysis of all cell types (**Fig. 2b**)), were subsampled randomly for the integrated data. Genotyping amplicon UMIs were downsampled (x100 iterations) to one per cell and mutant cell frequency was determined for each cluster for either the integrated dataset or individual samples. This frequency was then divided by the total mutant cell frequency across all progenitor subsets for each of the iterations. Linear mixed effects analysis was performed using the lme4 package (v.1.2-1). Progenitor identity was defined as the fixed effect, and for random effects, we used intercepts for individual patients (subjects) and iterative downsampling. P-values were obtained by likelihood ratio tests of the full model with the fixed effect against the model without the fixed effect^166^.

### RNA velocity

RNA velocity was calculated using scVelo (v0.2.2)^57^. For generating the loom file, the Python (v3.7) version of Velocyto (v0.17)^56^ was ran using the velocyto run command. The cell barcode and bam files were obtained using Cell Ranger. In addition to the cell barcode and bam files, a GTF file corresponding to the reference used for alignment (hg19; Ensembl 187) was supplied. Repetitive regions were masked using a GTF file downloaded from UCSC selecting for repetitive regions in GRCh37 (hg19). QC was assessed by the percent of unspliced reads per sample, requiring a minimum of 25% total unspliced reads. If duplicated gene names were present in the spliced and unspliced tables the counts were summed to leave only unique genes. Next, gene velocity for each patient and genotype was estimated separately using scVelo (v0.2.2). In order to avoid a potential confounder of unequal number of cells for each genotype, random sampling of the same number of mutant and wildtype cells to the minimum number in either group was performed for each patient sample for downstream analysis. Gene selection for RNA velocity estimation was performed requiring a minimum of 20 counts. After log-normalization by cell depth, the top 2,000 genes with the highest dispersion were selected for downstream calculations. Next, first and second order moments were computed among nearest neighbors in principal component space, using the pp.moments function with parameters n_pcs = 30 and n_neighbors = 30. RNA velocity was estimated using the dynamical model option of the tl.velocity function. The cell-to-cell transition probability matrices were retrieved for either wild type or mutant cells. For a given cell, we averaged the probabilities of transitioning to transcriptional states within a cluster of interest. This resulted in a mean probability of transition for the cell of interest to a given cluster. Statistical significance of the mean single cell differentiation probabilities between genotypes was estimated by linear mixed models. Sample was added as the random effect and genotype as the fixed effect. P-values were obtained by likelihood ratio tests of the full model with the fixed effect against the model without the fixed effect. To further compare wildtype to mutant probabilities for a given transition, we calculated the median of the distribution of single-cell mean transition probabilities toward other cell clusters, and calculated the mutant-to-wildtype odds ratio of the median probabilities.

### Gene module scoring, differential expression and gene set enrichment analysis

For examining gene and gene module expression (see **Supplementary Table 2**), the function AddModuleScore was used to calculate the relative expression of the genes for each cell within the Seurat package (e.g. **Fig. 2c**; MkP-EP module score (union of the MkP and EP module genes in **Supplementary Table 2**) was calculated using the AddModuleScore function)^44^. Briefly, control gene module expressions were calculated and subtracted from the average gene module expression of interest, as previously described^55^. All analyzed genes were classified based on average expression into 24 bins, and for each gene in the module, 100 control genes are randomly selected from the same expression bin as the gene of interest^55^. For statistical analysis, genotype status was entered as the fixed effect and subjects as random effects in a linear mixed model. P-values were obtained by likelihood ratio tests of the full model with the fixed effect against the model without the fixed effect.

Differential expression analysis comparing wildtype and mutant cells was conducted using a within-sample permutation test for each progenitor cell subtype. Briefly, to ensure equal representation from each patient, the numbers of mutated and wildtype cells from each patient were downsampled to the same number, respectively. Observed log2 fold change values were calculated with original genotyping assignments (MUT versus WT) for the tested genes. The tested genes included the top 2,500 most variable genes which were filtered for those expressed in at least 5% of either group (mutated versus wildtype), for each progenitor subtype. Ribosomal and mitochondrial genes were excluded. Next, over 100,000 iterations, WT and MUT labels were shuffled within each patient, and fold change values were re-calculated to create a background distribution. P-values were calculated per gene as a percent of permutations whose absolute fold change values were more extreme than the absolute value of the observed fold change (**Supplementary Table 3**). As an orthogonal approach, we also performed differential expression analysis comparing wildtype and mutant cells via the linear mixed model framework. For each gene, genotype status was entered as the fixed effect and subjects as random effects. P-values were obtained by likelihood ratio tests of the full model with the fixed effect against the model without the fixed effect (**Supplementary Table 3**).

Hypergeometric test for gene set enrichment analysis of the integrated differentially expressed genes (P-value < 0.05, log2(fold change) > 0.25) was performed using the Cluster Profile package (v. 0.1.9)^167^. FDR multiple hypothesis testing correction was performed. MSigDB C2: Chemical and genetic perturbations (CGP) sources were included in the analyses (**Supplementary Table 4**).

### Copy number variation analysis

The InferCNV package (v.1.4.0)^43^ was used to analyze the single cell dataset for any duplications or deletions of entire chromosomes or large chromosome fragments. Briefly, by comparing expression levels of genes annotated by chromosomal position (using the CONICSmat package, v0.0.0.1^168^) to a set of reference cells (in this case, a one-versus-rest comparison of cells by patient of origin), a heatmap of relative expression can be generated and used to identify regions with significantly increased or decreased expression. We removed the few genes for which alternative positions have been reported (<2% of genes). We downsampled our dataset to 978 genotyped cells from each patient (the minimum number of genotyped cells from any given individual patient). We then ran the InferCNV workflow with recommended parameters, using the i6 6-state Hidden Markov model (**Extended Data Fig. 2a**). As a positive control, we specifically analyzed relative expression of Y-chromosome genes to ensure sex-differences between patients were appropriately reflected in our data (**Extended Data Fig. 2b**).

### Hypomethylated motif enrichment analysis in differentially expressed genes

The HOMER (v4.9) scanMotifGenomeWide function was used to search for occurrences of the *DNMT3A* R882 hypomethylated motif and a control motif containing a CpG. For each gene in the scRNA-seq dataset, TSS coordinates were identified and a .bed file was created with intervals of ±10 kb, 30 kb or 50 kb surrounding each TSS. These two sets of coordinates were intersected using bedtools (v2.30.0), and the number of hypomethylated motif or control motif sites were counted per gene. Differentially expressed genes were classified as upregulated (P < 0.05, log2(fold change) > 0.25) or downregulated (P < 0.05, log2(fold change) < − 0.25), and counts of hypomethylated motif sites were compared, with P-values obtained by Wilcoxon rank sum test. To ensure that the results were not driven simply by the presence of a CpG, we also determined the ratio of the counts of the hypomethylated motif to that of the control shuffled motif with CpG per gene.

### Joint multiplexed single-cell methylome and single-cell RNA-seq library construction

DNA methylation data was processed produced as previously described by Gaiti et al.^39^ Briefly, genomic DNA (gDNA) and mRNA were separated as follows. A modified oligo-dT primer (5′-biotin-triethyleneglycol-AAGCAGTGGTATCAACGCAGAGTACT30VN-3′, where V is either A, C or G, and N is any base; IDT) was conjugated to streptavidin-coupled magnetic beads (Dynabeads, Life Technologies) according to the manufacturer’s instructions. To capture polyadenylated mRNA, we added the conjugated beads (10 μl) directly to the cell lysate and incubated them for 20 min at room temperature with mixing to prevent the beads from settling. The mRNA was then collected to the side of the well using a magnet, and the supernatant, containing the gDNA, was transferred to a fresh plate. Single-cell complementary DNA was amplified from the tubes containing the captured mRNA according to a variation of the Smart-Seq2 protocol ^107^ using molecular crowding to increase sensitivity^169^. After amplification and purification using 0.8X SPRI beads, 0.5 ng cDNA was used for Nextera Tagmentation and library construction. At the cDNA amplification step, the following primers were spiked-in (0.5 μM final) to specifically increase capture of the locus around *DNMT3A* R882 mutation (Fw: 5’- TCGTCGGCAGCGTCAGATGTGTATAAGAGACAGGGTTTCCCAGTCCACTATACTGACG-3’ ; Rv: 5’- TCGTCGGCAGCGTCAGATGTGTATAAGAGACAGATGACCGGCCCAGCAGTCTC -3’). The same primers were used to specifically amplify the target locus separately in a portion of the cDNA. Library quality and quantity were assessed using Agilent Bioanalyzer 2100 and Qubit, respectively. Libraries were then sequenced with paired-end, 50-base reads, using a NovaSeq sequencer (Illumina).

Genomic DNA present in the pooled supernatant and wash buffer from the mRNA isolation step was concentrated on 0.8X SPRI beads and eluted directly into the reaction mixtures for single digest or Msp1 + HaeIII (Fermentas) for double restriction enzyme digest reaction (10µL final reaction) for 90 min at 37°C. Heat-inactivation was performed for 10 min at 70°C. Digested DNA was filled-in and A-tailed at the 3’ sticky ends in 8.5 μL final volume of 1X CutSmart with 2.5 units of Klenow fragment (Exo-, Fermentas). Reaction was supplemented with 1 mM dATP and 0.1 mM dCTP and 0.1 mM dGTP (NEB) and performed as follows in a thermocycler: 30°C for 25 min, 37°C for 25 min, and 70°C for 10 min (heat-inactivation). Custom barcoded methylated adaptors (0.1 μM) were then ligated overnight at 16°C with the dA-tailed DNA fragments in the presence of 800 units of T4 DNA ligase (NEB) and 1 mM ATP (Roche) in a final volume of 11.5 μL of 1X CutSmart buffer. T4 DNA ligase heat-inactivation was performed at 70°C for 15 min the next day. Genomic DNA from 24 individual cells were pooled together according to their barcodes, giving 4 pools of 24 cells for a 96-well plate. Pooled genomic DNA was cleaned-up and concentrated using 1.8X SPRI beads (Agencourt AMPure XP, Beckman Coulter). Each pool was then converted using an enzyme-based conversion to increase the recovery of single cell gDNA compared to standard bisulfite conversion (NEBNext Enzymatic Methyl-seq, New England Biolabs)^102^. Standard bisulfite conversion was implemented for double restriction enzyme digest reactions, as previously described^107^. Converted DNA was then amplified using primers containing Illumina i7 and i5 index. Following Illumina pooling guidelines, a different i7 and i5 index was used for every 24-cell pool, allowing multiplexing of several samples for sequencing on Illumina NovaSeq6000. Library enrichment was done using KAPA HiFi Uracil+ master mix (Kapa Biosystems) and the following PCR condition was used: 98°C for 45 secs; 6 cycles of: 98°C for 20 secs, 58°C for 30 secs, 72°C for 1 min; followed by 12 cycles of: 98°C for 20 secs, 65°C for 30 secs, 72°C for 1 min. PCR was terminated by an incubation at 72°C for 5 min. Enriched libraries were cleaned-up and concentrated using 1.3X SPRI beads. DNA fragments between 200 bp and 1 Kb were size-selected and recovered after resolving on an E-Gel EX Precast Agarose Gels (Thermo Fisher Scientific). Library molarity concentration calculation was obtained by measuring concentration of double stranded DNA (Qubit) and quantifying the average library size (bp) using an Agilent Bioanalyzer. Every 24-cell pool was mixed with the other pools in an equimolar ratio. Negative controls (empty wells with no cells) were used to control for non-specific amplification of the libraries.

### Multimodal single cell methylome and RNA sequencing data processing

#### Methylation analysis pipeline

DNA methylation data was processed as previously described^39^. Pools of 24 cells were demultiplexed based on a supplied list of cell barcodes. Adapter sequences were trimmed by the first 3 bp on each 3’ end of R1 and R2. Bismark (v0.14.5) was used to create bisulfite-converted genomes of GRCh38 (hg38 Ensembl version 93). Reads were mapped using Bismark with Bowtie (v2.2.8) and default alignment parameters. BAM files were then used to run Bismark methylation extractor ignoring 6 bp from the end of R1 and 5 bp from R2. This was done to remove technical variability introduced at the ends of the reads during end repair with unmethylated nucleotides. These settings were determined from the M-bias reports, which contain the methylation proportion at each read position. Bismark methylation extractor (-bedgraph comprehensive) was used to determine the methylation state of each individual CpG. Cells with > 99% conversion efficiency as determined by Bismark were retained for downstream analysis. Reads mapping to ChrY and the mitochondrial genome were removed from the resulting .cov files. For all downstream analysis, the methylation status of CpGs per cell was binarized. CpGs with 10-90% methylation values were removed (< 2% of total CpGs) and those with values <10% were encoded as 0, while those with values >90% are encoded as 1. On average, 209,519 ± 15,200 (± SEM) unique CpGs per cell were covered in the DNA methylome.

#### RNA analysis pipeline

scRNA-seq data was aligned using STAR (v2.5.2a). Default parameters were used, other than twopassMode Basic. Reads were aligned to GRCh38 (hg38 Ensembl version 93). Gene counts were determined using featureCounts from Subread (v1.5.2) using default parameters. Ensembl gene IDs were converted to hgnc symbols using the R package biomaRt (v2.40.5). In cases where there were duplicated gene symbols the counts were summed. Seurat (v3.1.1) was then used to analyze gene expression data. Cells were filtered for mitochondrial reads of less than 25% and a minimum of 200 detected genes. Genes were filtered for coverage across at least three cells. The mean (± standard deviation) number of detected genes was 5,763 ± 2075 genes/cell (range 3,117 ± 678 – 8,715 ± 1,449 genes/cell across the plates). The mean number of reads was 511,840 ± 315,941 reads/cell (range 170,383 ± 63,951 – 779,771 ± 361,887 reads/cell across the plates). Normalization and variable feature detection were performed for each batch (i.e. plate). Batch correction and integration was performed via the Seurat integration pipeline^44^ using recommended parameters for SelectIntegrationFeatures, FindIntegrationAnchors, and IntegrateData. Dimensionality reduction was performed by principal component analysis using the RunPCA function, and the first 12 principal components were retained for downstream analysis. For visualization, UMAP^164^ was performed using the RunUMAP function. Cell type assignment was performed as described for the 10x Genomics scRNA-seq data.

#### Genotyping

To process genotyping data, genotyping FASTQs were aligned the same manner as RNA library FASTQs. Pysam (v0.8.2.1) was used to select reads overlapping the target allele by using the pileup function. Reads were filtered by a minimum read mapping quality (MAPQ) of 40 and a minimum base quality (Phred score) of 20. Each remaining read was classified as either mutant or wildtype based on the nucleotide detected at the mutation site based on bulk sequencing data^40^. Cells were classified as mutant if there were at least two mutant reads, and wildtype if there were at least three wildtype reads (increased stringency given mutation heterozygosity) and no mutant reads. For genotyping libraries with increased sequencing depth (7,712 ± 319 versus 20 ± 2.75 reads; mean ± SEM), the base quality thresholds were increased to 40. For genotype classification, a bootstrapping approach was implemented by randomly sampling 50 reads for 100 iterations. For each iteration, a mutant fraction cutoff of 0.10 was applied. The final genotyping call was performed in cells with above 80% bootstrap support.

### Average Single Cell Methylation

We compared single cell methylation at selected genomic regions (i.e. enhancers, CpG islands, ChIP-seq peaks) between mutant and wildtype cells from each patient. To achieve this, we first filtered for CpG sites with coverage in at least three cells in each patient, in order to reduce inter-patient variability. The genomic region of interest was then intersected with the CpG sites using the R package GenomicRanges (v1.36.1). Finally, the average methylation for a given region across the covered CpG sites was calculated for each cell. Statistical significance between genotypes was estimated by linear mixed models. Sample was added as the random effect and genotype as the fixed effect. P-values were obtained by likelihood ratio tests of the full model with the fixed effect against the model without the fixed effect. Due to potential differences between single versus double digest data, we display single digest datasets as representatives (unless otherwise indicated for analysis that specifically relies on the enhanced coverage of double digest).

### Single-cell differentially methylated region (DMR) analysis to identify preferential hypomethylation

For each cell, Bismark methylation extractor output files (containing information on methylation state of each individual CpG) were intersected with the genomic regions of interest (*e.g.*, promoters) using BEDTools (v2.27.1). A generalized linear model (GLM) was then built to predict the DNAme for a given genomic region between genotypes, accounting for global methylation changes. For each cell, the global DNAme value was defined as the average DNAme across all genomic regions investigated. The model used was as follows:

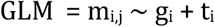

Where m_i,j_ represents the average DNAme of the genomic region j (*e.g*., promoter of *FOXA2*) for cell i; gi represents the genotype of cell i and ti represents the average methylation for all CpGs detected in cell i. Only genomic regions with sufficient DNAme information (>5 CpGs per region for promoters and >50 CpGs for ChIP-Seq peaks) in at least 15 cells per group (mutated or wildtype) were used in the analysis. To test the impact of genotype on DNAme for a given genomic region (*e.g*. promoter of *FOXA2*), P-values were derived from the GLM (calculated from the t-statistic computed by dividing the genotype (g) regression coefficient by the residual standard error, **Supplementary Table 5**). To calculate the percentage methylation difference in mutant cells for a given genomic region of interest, the average across mutant and wildtype cells was taken within plate to control for batch effects. Next, the DNAme difference between mutant and wildtype was computed within plate and a weighted average of the difference was calculated, using the number of cells from each plate as weights. In order to be consistent across genes, promoters were defined as 1 kb upstream and 1 kb downstream of transcription start sites (hg38 RefSeqGene)^109^. ChIP-seq peaks were obtained from ENCODE (hg38 Tfbs clustered)^109^. When directly examining the methylation status of SUZ12 and EZH2 targets, we intersected the ENCODE ChIP-seq peaks with bivalent peaks (H3K27me3, H3K4me3) from human CD34^+^ hematopoietic progenitor cells^110^.

#### Gene set enrichment analysis

To define the pathways enriched at hypo- or hypermethylated TSS, genes were ranked based on methylation difference, and differentially hypomethylated genes (P < 0.05) were selected as inquiry for pathway analysis. We note that gene set enrichment analysis of RRBS data may be confounded by the fact that the use of restriction enzymes enriches for CpG rich genomic regions as well as CpG rich promoters. Thus, pathway enrichment was performed via a pre-ranked gene set enrichment approach (and thus including only genes covered in our data) using the msigdbr (v7.2.1) and fgsea (v1.12.0) R packages, with the MSigDB C2 CGP collection of curated gene sets.

### *DNMT3A* R882 motif analysis

#### CpG flanking motif analysis

To identify the sequences surrounding hypo or hypermethylated CpG sites in wildtype versus *DNMT3A* mutant hematopoietic progenitors, we first performed differentially methylated regions (DMR) analysis in CpG islands as described above in the “Single-cell differentially methylated region (DMR) analysis” section. CpGs within hypo or hypermethylated regions (P < 0.05) were selected, and the surrounding ± 6 bp sequences were extracted using bedtools (v2.25.0). The frequency of each base pair at each position relative to the CpG site was calculated, and statistical significance was assessed by Fisher exact test. Odds ratio logo was generated by calculating the frequency for each base at each position for either hypomethylated or hypermethylated CpG sites. To identify differentially enriched bases surrounding the CpG site, we applied increasingly stringent thresholds on the absolute methylation difference required between wildtype and mutated cells to consider the sites, and estimated the odds ratio of base frequency of hypo-over hyper-methylated sites at a given position relative to the CpG site. Next, we calculated the correlation between the methylation difference required and the odds ratio of base frequency. We define bases differentially enriched or depleted in hypo-versus hyper-methylated based on the correlation significance (P < 0.05). For CpG sites with greater than absolute methylation difference of 0.5, the odds ratios were computed and used as input to generate the logo using the ggseqlogo (v0.1) package. To identify transcription factors with the motif pattern of interest, we used the HOCOMOCO v11 human motif position weight matrix (PWM) collection in HOMER format with *P* < 0.001. For each of the PWMs, we selected the position containing the highest CpG probability and calculated the similarity score of the flanking −2 and +2 positions relative to the CpG site against the hypo-methylated flanking sequences, based on the correlation of the base frequencies along each of the motifs.

### Average methylation at MYC motifs and modeling regulon expression

The MYC and ARNT motif PWM was downloaded from the HOCOMOCO (v11) human TF database and used as input to HOMER (v4.9). The scanMotifGenomeWide function was used to search for occurrences of motifs throughout the genome. The R package GenomicRanges (v1.36.1) was used to intersect CpG sites with motifs and respective ChIP-seq peaks (ENCODE database)^109^. Methylation per cell was then averaged across the covered CpG sites. Positively regulated downstream MYC targets were determined using pySCENIC (v0.10.0). Counts were converted to transcripts per million (TPM) and genes in the count matrices were filtered for those in the cisTarget database (all available hg38 files were used). The hgnc (v9) motif file from the cisTarget database was used to generate a list of input motifs. Regulons were determined from each patient sample separately with default parameters as described^170^. To analyze expression of the regulons, per-cell AUC scoring was done using the aucell function. The relationship between MYC motif methylation and regulon expression was modeled with a generalized linear model (GLM) using a Gamma distribution with the following model:

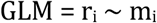

Where r_i_ represents the AUC score to MYC downstream targets for cell i; mi represents the DNAme of MYC motifs for cell i. Due to batch effects between methylome sequencing methods, only samples that were prepared using the enzymatic method were included. Rare outliers were excluded that had a Cook’s distance greater than 2 * mean Cook’s distance. To test the impact of MYC motif methylation on regulon expression, the P-value was derived from GLM output (calculated from the t-statistic computed by dividing the MYC motif methylation (m) regression coefficient by the residual standard error).

### AML PRC2 target methylation analysis

Methylated base call files of *DNMT3A*-mutated AML samples were downloaded from Glass et al.^105^ PRC2 targets were obtained from the union of EZH2 and SUZ12 ChIP-seq peaks (see “single-cell differential methylation analysis”), as approximately 50% of SUZ12 ChIP-seq peaks overlapped with EZH2 peaks. PRC2 targets were further intersected with promoters using the GenomicRanges (v1.38.0) findOverlaps function, requiring at least 30 bp to be overlapping. We note that PRC2 targets are known to have a higher CpG content^171,172^, potentially biasing the result given the higher coverage of RRBS of high CpG content promoters. We therefore also compared PRC2 target methylation only with high CpG content promoters as annotated by Saxonov et al.^173^ and ± 1 KB surrounding the TSS. For each sample 270,000 CpG sites were randomly sampled from either promoters overlapping with PRC2 peaks, or non-overlapping promoters as a control. The number of randomly sampled CpG sites was selected based on the minimum coverage among replicates. The ratio of methylation between *DNMT3A* mutant and wildtype AML (**Fig. 4h**), required to pair each mutated AML with a wildtype sample. As this pairing is arbitrary (i.e., samples are not explicitly matched), to safeguard against a non-representative pairing, we permutated all possible pairing and P-values were obtained by Wilcoxon rank sum test. The example shown represents the median P-value among the permutations. Methylated base call files of *DNMT3A*-mutated and wildtype AML samples were downloaded from TCGA^118^. Overlap of PRC2 ChIP-seq peaks and promoter regions was carried out as described above. The average methylation at high CpG promoters that overlap with PRC2 peaks and high CpG promoters that do not overlap with PRC2 peaks was calculated per sample and compared between *DNMT3A* R882 mutant and *DNMT3A* wildtype AML (Wilcoxon rank sum test).

### Single nucleus ATAC-sequencing of Dnmt3a R878 and wildtype HSPCs

Hematopoietic progenitors (Lin-1^−^, c-Kit^+^) were sorted from wildtype (n = 3 mice) or *Dnmt3a* R878H (n = 3 mice) via c-Kit enrichment as directed by the manufacturer (CD117 Microbeads, clone 3C1, Miltenyi, Auburn, CA; LS Columns (Cat. No. #130-042-401), Miltenyi) followed by FACS (Lin-1 BV421 (Cat. No. #133311), Biolegend, San Diego, CA; CD117 APC (clone 2B8, Invitrogen, Waltham, MA). Nuclei isolation was performed as suggested by the manufacturer (10x Genomics, Pleasanton, CA). Briefly, single cell suspensions were centrifuged at 300 rcf for 5 minutes and cell pellets were resuspended in 100 μl of lysis buffer (Tris-HCl pH 7.4, 10mM; NaCl 10mM; MgCl2 3mM; Tween-20 0.1%; Nonidet P40 substitute (Sigma-Aldrich, St. Louis, MO) 0.1%; Digitonin 0.01%; BSA 1%; DTT 1 mM; RNase inhibitor 1 U/μL (Sigma-Aldrich, St. Louis, MO)) and kept on ice for 3 minutes. Then, 1 ml of wash buffer (Tris-HCl pH 7.4, 10mM; NaCl 10mM; MgCl2 3mM; BSA 1%; Tween-20 0.1%; DTT 1 mM; Sigma Protector RNase inhibitor 1 U/μL) was added. The isolated nuclei were centrifuged for 5 min at 500 rcf, and pellets were resuspended in Diluted Nuclei Buffer (10x Genomics Nuclei Buffer 1X; DTT 1 mM; Sigma Protector RNase inhibitor 1 U/μL). Nuclei concentration was determined by hemocytometer and processed as indicated by the manufacturer (10x Genomics User Guide: Chromium Next GEM Single Cell Multiome ATAC + Gene Expression, CG000338). Single nucleus ATAC and Gene Expression (GEX) libraries were constructed in parallel and assessed for quality control metrics using Agilent Bioanalyzer 2100 and Qubit respectively. ATAC libraries were sequenced to a depth of 25,000 read pairs per nucleus (paired-end, dual indexing: Read 1N 50 cycles, i7 Index 8 cycles, i5 Index 24 cycles, Read 2N 49 cycles) and GEX libraries were sequenced to a depth of 20,000 read pairs per nucleus (paired-end, dual indexing: 28 cycles for Read 1, 10 cycles for i7 Index, 10 cycles for i5 Index, 90 cycles for Read 2).

### Single nucleus ATAC-sequencing data processing

Pre-processing was performed using 10x Genomics Cell Ranger ARC (v1.0.1). Reads were de-multiplexed using the cellranger-arc mkfastq function. Single cell feature counts for each sample were then generated using the cellranger-arc count function. The gene expression information for these libraries exhibited exceedingly low UMI and genes per cell consistent with lower quality RNA in single-cell nuclei Multiome data; as such, we moved forward utilizing only the ATAC data for analysis. ATAC data was processed using the ArchR package (v1.0.1) ^174^ using the atac_fragments.tsv.gz file generated by the cellranger-arc count function as input. Arrow files were created using a minimum TSS enrichment score of 5 and a minimum number of unique nuclear fragments of 1,000. Doublet scores were calculated using the addDoubletScores function with k = 10, knnMethod = “umap” and LSImethod = 1. Doublets were removed using the filterDoublets function with default parameters. Dimensionality reduction was performed through iterative semantic index (LSI) using the cell by genomic window (500 bp) matrix as input, using the addIterativeLSI function with the following parameters: iterations = 3, resolution = 0.2, sampleCells = 1,000, var.features = 25,000 and dimsToUse = 1:30. Cell clusters were identified using the addClusters function using the iterative latent semantic index (LSI) dimensions as input, with method = “Seurat”, resolution = 0.8. For visualization, UMAP dimensionality reduction was performed using the LSI dimensions as input, using the addUMAP function with: nNeighbors = 30, minDist = 0.5 and metric = “cosine”. Cell identities were assigned based on gene accessibility scores of known marker genes. Custom motif accessibility deviations were calculated as follows: position weight matrices in HOMER format (*P* < 0.001) were downloaded from the HOCOMOCO v11 mouse database. Motif occurrences were identified using the scanGenomeWide function of the HOMER package. To include only high confidence motif sites, we applied a minimum odds ratio score threshold of 6. We next created custom peakAnnotations using ArchR and performed ChromaVar analysis using the addDeviationsMatrix function with default parameters.

### CH05 sample processing and analysis

#### Single cell RNA-seq processing and downstream analysis

CH05 bone marrow underwent sorting, scRNA-sequencing and genotyping with GoT as described above for samples CH01-04, with the exception of the addition of the CITE-seq integration. Briefly, the Total-seqA antibodies (Biolegend: CD38, CD9, CD49f, CD45RA, CD41, CD36, CD69, CD42, CD14, CD71, CD45RB, CD45RO, CD37, CD7, CD279, CD47, CD90, CD99, CD84, CD274, FLT3, CD79B, CD45, CD81) were used according to manufacturer’s recommendations. The CD34^+^ sorted cells were incubated with the antibodies for 30 minutes and underwent washes 3X. 10x data were processed using Cell Ranger (v3.0.1) with default parameters. Reads were aligned to the human reference sequence hg19. Control bone marrow samples (BM01-05) were identified from previously published reports^141,142^ with raw count matrices available for download. The Seurat package (v.3.1) was used to perform integration and unbiased clustering of the CD34^+^ sorted cells from patient samples as described previously with the following notable exceptions^163^. The publicly available archived count matrices for samples BM04 and BM05 had the following QC filtering: the mitochondrial and ribosomal genes were removed, and only cells with > 400 unique genes and between 1,000 and 10,000 UMIs were kept. Consequently, these two patients were not filtered with the aforementioned criteria. CH05 and BM01-03 were filtered identically as samples CH01-04, following which mitochondrial and ribosomal genes were removed from the gene expression matrix. All samples were then normalized and integrated as described previously, with the exception of proportion of mitochondrial genes no longer being regressed out as a potential confounder. We identified 26 clusters in the integrated data, which were annotated as above using lineage markers previously identified for normal hematopoietic progenitors^53,175^.

Following cell-type assignment, we down-sampled the count matrices using the downsampleBatches function from the scuttle package (v1.0.4) to ensure that the average per-cell geometric mean of raw counts was consistent across all 6 patient samples^176^.

Module scores were calculated as described above. The performance of the CITE-seq antibodies was assessed based on expected expression patterns across the progenitor subsets.

### Single nucleus ATAC-seq and downstream analysis

snATAC-seq data for CH05 was generated as described above using the Multiome platform (10x Genomics) and GoT performed as described above using the cDNA generated from the Multiome workflow. The gene expression information for these libraries exhibited very low UMI and genes per cell consistent with lower quality RNA in single-cell nuclei Multiome data; as such, we moved forward utilizing only the ATAC data for analysis. For the analysis, fragment files were generated by processing the fastq files using cell-ranger-ARC (v.1.0.0). Downstream analysis was performed using the ArchR (v1.0.1) pipeline^174^. Based on the distribution of total fragments and TSS enrichment per cell, empty droplets were filtered out by requiring a minimum of 3,000 fragments per cell and a TSS enrichment score of 7.5. Potential doublets were detected using the addDoubletScores function, using KNN on the UMAP dimensionality reduction with k = 10. Cell barcodes with high enrichment for doublet scores were removed using the filterDoublets function with default parameters. Next, we performed dimensionality reduction through iterative latent semantic indexing (LSI) using the top 25,000 variable features. Cell clustering was performed using the addClusters function, with the following parameters: reduceDims = “IterativeLSI”; method = “Seurat”; resolution =1. For visualization, further dimensionality reduction was performed by applying UMAP to the iterative LSI space using the addUMAP function with the following parameters: nNeighbors = 30; minDist = 0.5; metric = “cosine”. Cell type identification was performed by manually inspecting the genes showing up-regulated gene accessibility scores (FDR < 0.01 and log2(fold change) > 1.25) for each of the defined clusters (**Extended Data Fig. 13c**). Motif occurrences were defined using the position weight matrices (PWMs) obtained from the Hocomoco (v.11.0) motif database or our custom PWMs for hypo-methylated and shuffled motifs using HOMER (v4.9), requiring a minimum enrichment score above 6. Transcription factor, hypo-methylated and shuffled motif accessibility was calculated using ChromVAR^177^ within the ArchR (v1.0.1) pipeline^174^. Supervised pseudotime trajectories for either erythroid or lymphoid fates were defined within the ArchR (v1.0.1) pipeline^174^ applying the addTrajectory function.

---

**Extended Data Figure 1.**
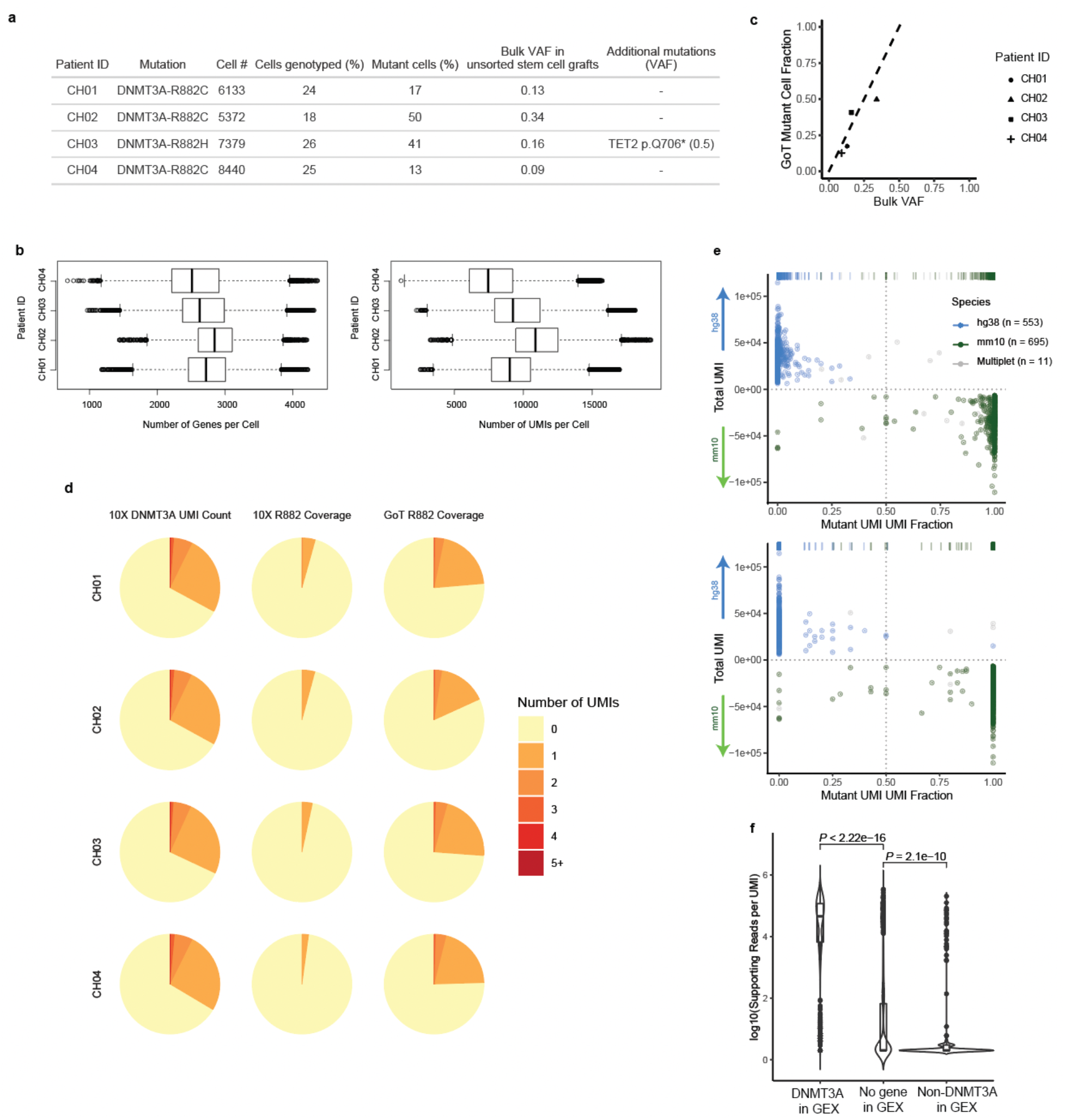
GoT captures genotyping information of thousands of CD34^+^ cells in scRNA-seq. **a,** Summary of GoT data from CH patient samples with *DNMT3A* R882 mutations. **b,** Number of genes per cell (left) and number of UMIs per cell (right) from CD34^+^ sorted hematopoietic progenitors by patient sample after QC filters. **c,***DNMT3A* R882 mutant fraction of single cells determined by GoT versus *DNMT3A* R882 mutation variant allele frequencies (VAF) in bulk sequencing of matched unsorted stem cell product. **d,** Fraction of cells by number of *DNMT3A* UMIs in standard 10x Genomics data without genotyping information (left), *DNMT3A* UMIs with R882 locus coverage in standard 10x data (middle), and *DNMT3A* UMIs with R882 locus coverage in GoT amplicon library (right). **e,** Species-mixing experiment data in which mouse cells (Ba/F3) with a human mutant *CALR* transgene were mixed with human cells (UT-7) with a human wildtype *CALR* transgene. Mouse and human genome alignment of 10x data with genotyping data from GoT pre (top) and post (bottom) implementation of UMI consensus assembly based on Levenshtein distance (online methods). **f**, Number of duplicate reads supporting cell barcode-UMI pair in the GoT library that is identified in the 10x gene expression (GEX) library as a *DNMT3A* gene (left), no gene (middle), or a non-*DNMT3A* gene (right).

**Extended Data Figure 2.**
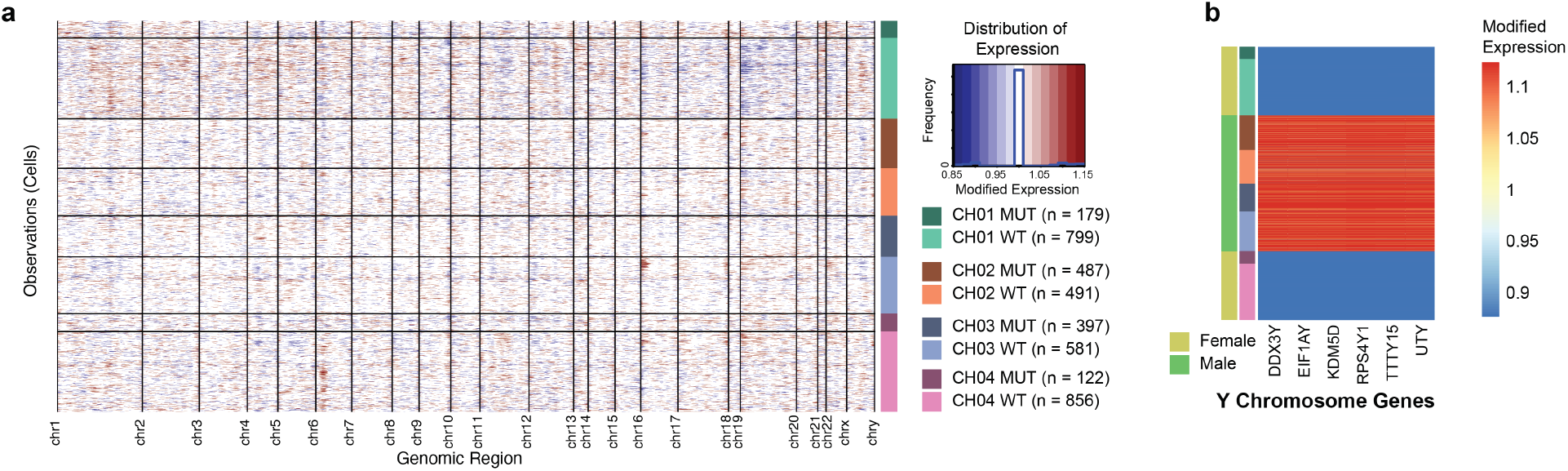
Copy number analysis of wildtype and mutant single cells from clonal hematopoiesis patient samples with*DNMT3A* sR882 mutations. **a,** Heatmap of relative expression of genes ordered by chromosome/chromosomal position following copy number variation analysis using the InferCNV package. Cells (y-axis) are stratified by patient and *DNMT3A* R882 genotype status. **b,** Heatmap of relative expression of Y-chromosome genes following copy number variation analysis and cell stratification as in **a**.

**Extended Data Figure 3.**
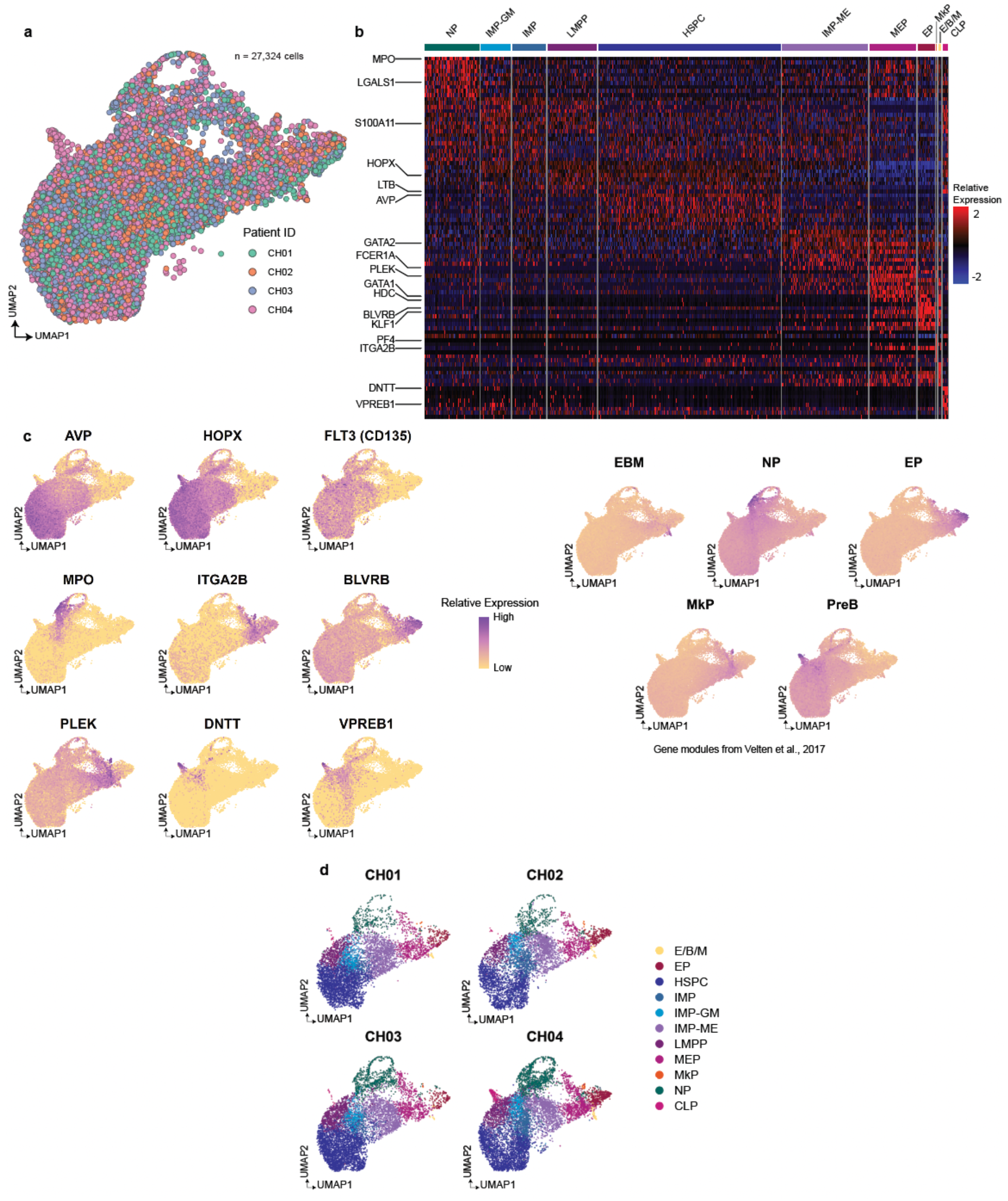
Integration of *DNMT3A* R882 mutation and assignment of progenitor subsets in clonal hematopoiesis patient samples. **a,** UMAP of CD34^+^ progenitor cells from samples CH01-CH04 after integration using the Seurat package (online methods). **b,** Heatmap of top 10 differentially expressed genes for progenitor subsets. **c,** Lineage-specific genes (left) and modules from Velten et al. (right, **Supplementary Table 2**) are scored and projected onto the UMAP representation of CD34^+^ cells. **d,** UMAP of CD34^+^ cells overlaid with cluster assignments, split by patient sample.

**Extended Data Figure 4.**
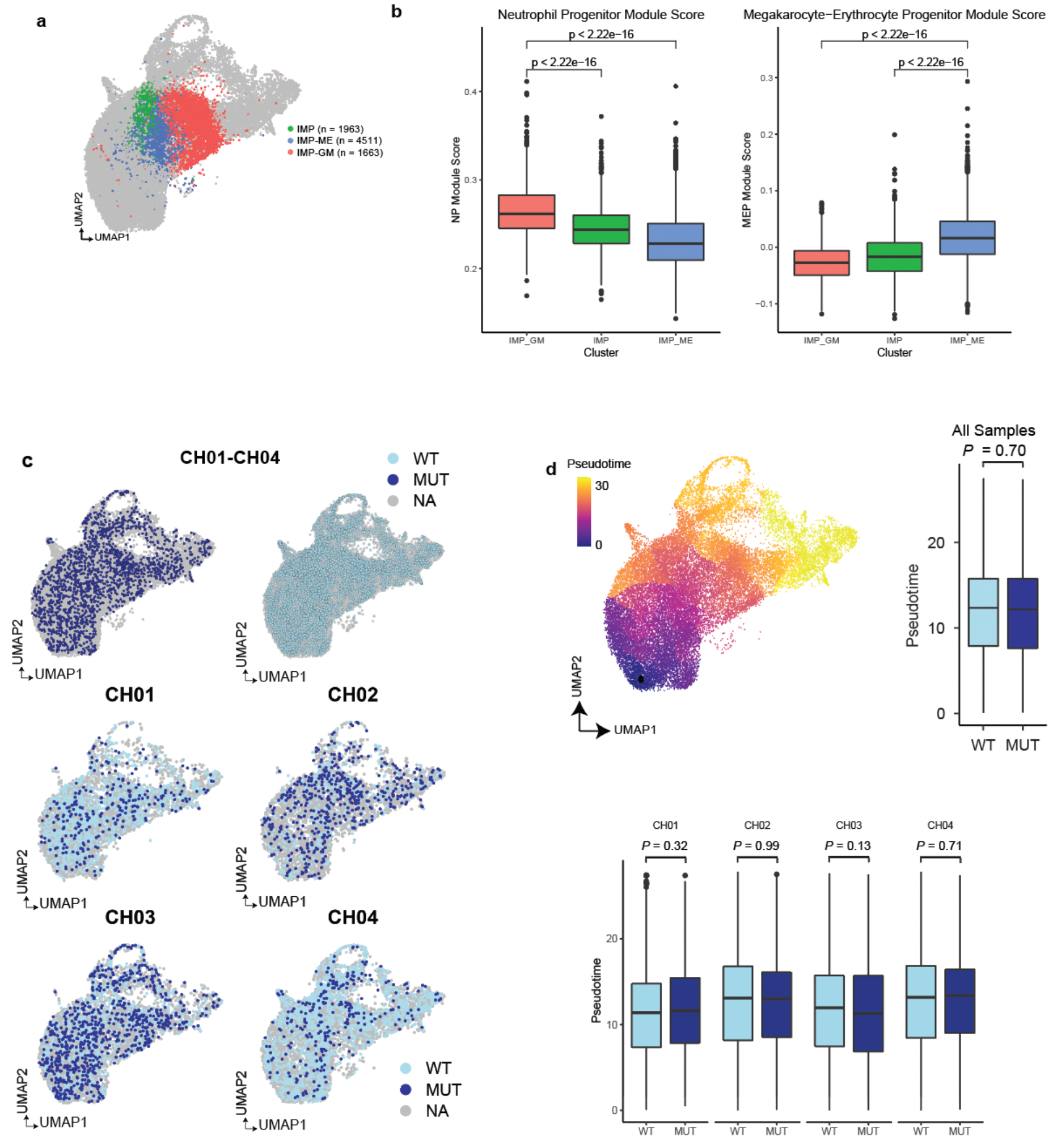
Classification of IMPs showing lineage biases and pseudotime analysis between mutated and wildtype cells. **a,** UMAP of CD34^+^ cells, overlaid with cluster assignment of all IMP subsets in the dataset. **b,** Neutrophil and Megakaryocytic-Erythroid lineage specific gene module scores from Velten et al. compared across the three IMP clusters. P-value was calculated from Wilcoxon rank sum test. **c,** UMAP of CD34^+^ cells overlaid with mutation status for WT, *DNMT3A* R882 mutant (MUT), or unassigned (NA), split by genotype for all samples (top) and by patient sample (bottom). **d,** UMAP with projected pseudotime values (top left). Pseudotime comparison between WT and MUT cells for all samples (top right) and for individual samples (bottom) as estimated by Monocle. P-value was calculated from likelihood ratio test of linear mixed model with/without mutation status for aggregate analysis (online methods, top) and Wilcoxon rank sum test for individual samples (bottom).

**Extended Data Figure 5.**
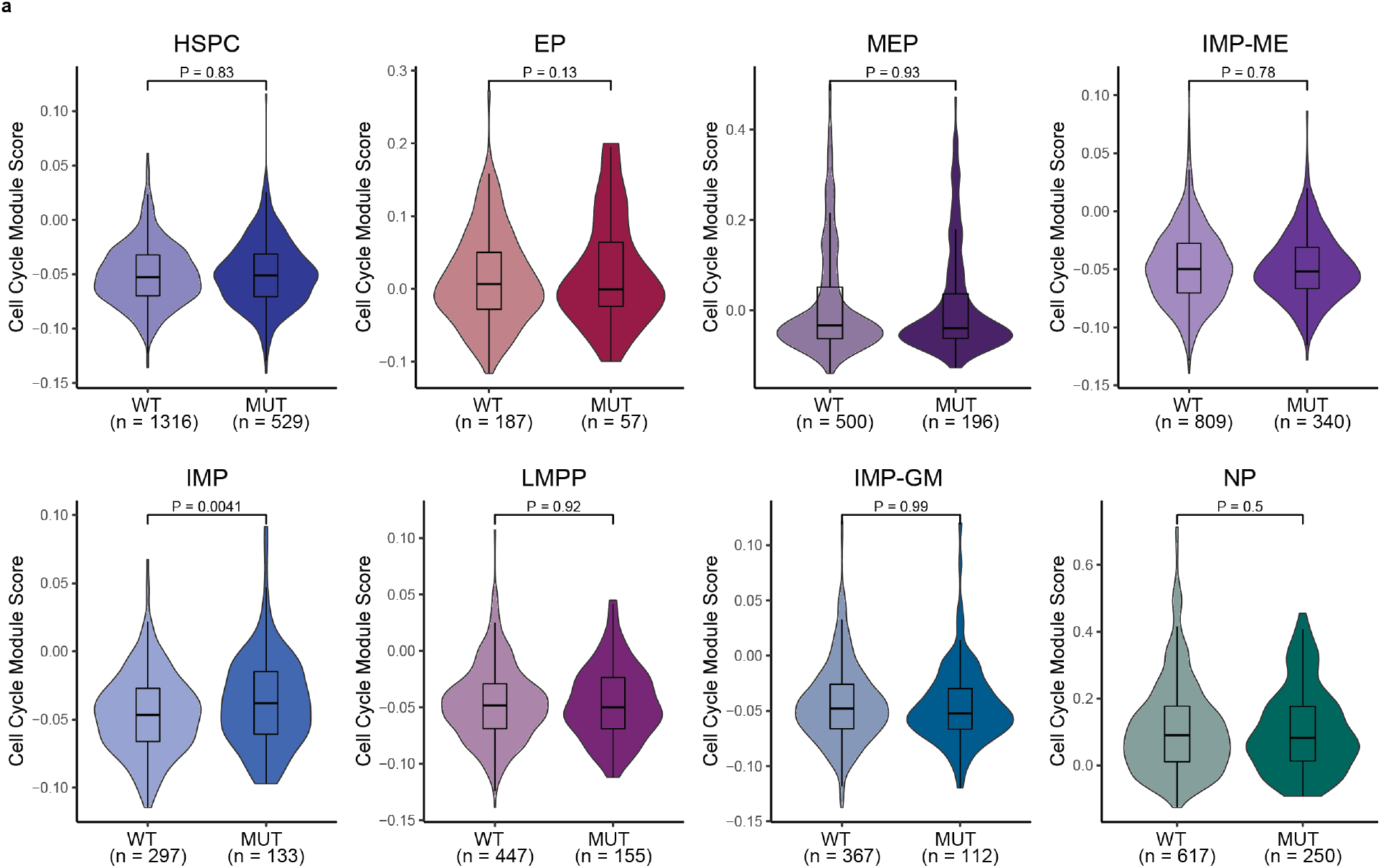
Cell cycle module expression comparison between mutated and wildtype progenitor cells. **a,** Cell cycle module score represents the union of S-phase and G2M-phase gene-module expression (**Supplementary Table 2**). P-value was calculated from likelihood ratio test of linear mixed model with/without mutation status (online methods). Analysis was performed for clusters with at least 200 genotyped cells across all patient samples.

**Extended Data Figure 6.**
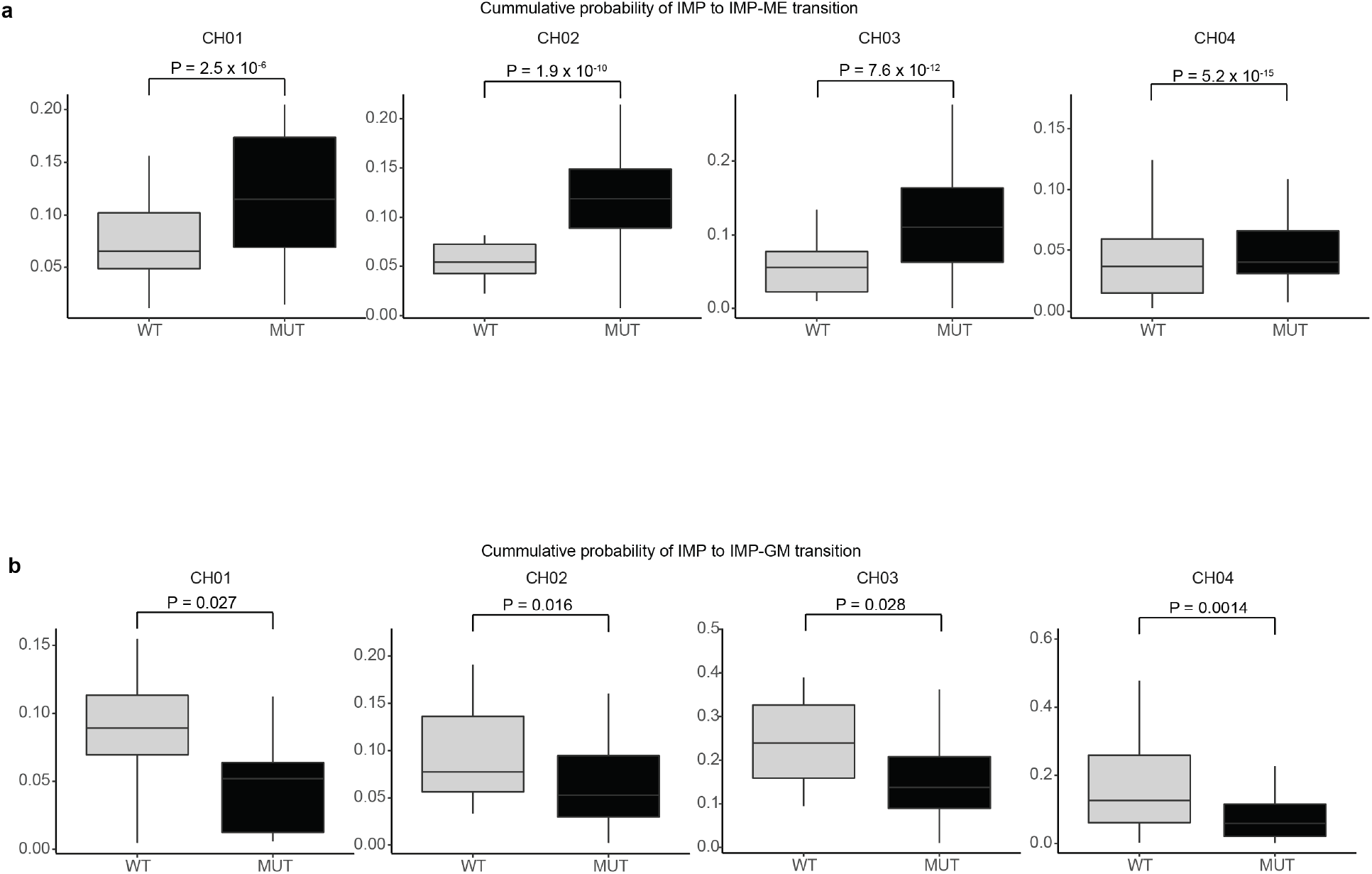
Transition probabilities via RNA velocity reveals a megakaryocytic-erythroid bias of IMPs. **a,** Single cell mean IMP → IMP-ME and **b,** IMP → IMP-GM transition probabilities, as measured via RNA velocity, between wildtype or *DNMT3A* R882 mutant IMPs for each sample. P-values from Wilcoxon rank-sum test.

**Extended Data Figure 7.**
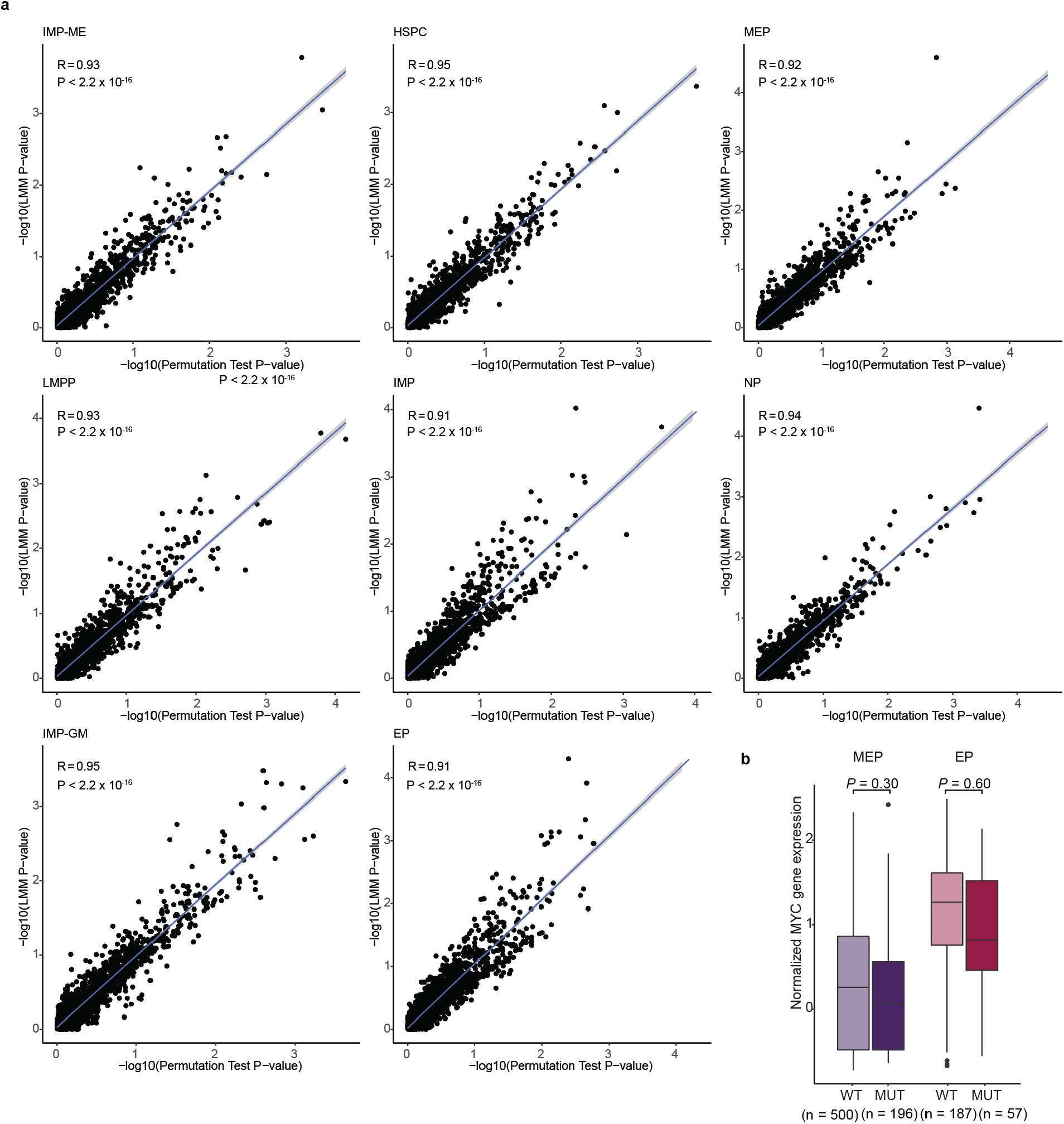
Comparison of differential expression analysis between permutation test and linear mixed model and MYC gene expression. **a,** P-values from permutation test and linear mixed model (online methods) are plotted per gene. Correlation coefficient R calculated using Pearson’s Correlation. P-values derived from Student’s t-distribution. **b,** Normalized *MYC* gene expression between mutated and wildtype cells in MEP and EP. P-value was calculated from likelihood ratio test of linear mixed model with/without mutation status (online methods).

**Extended Data Figure 8.**
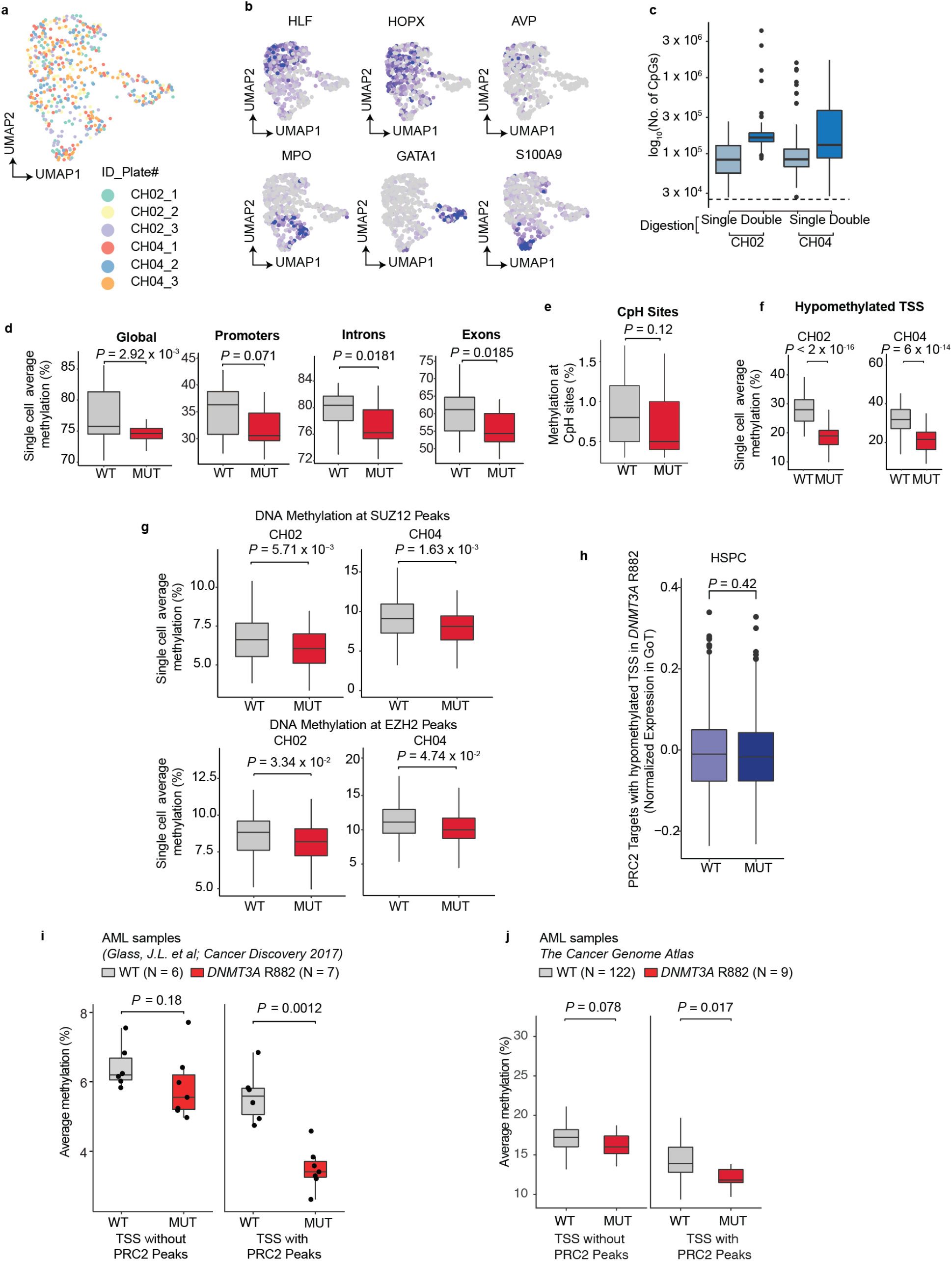
Multi-omics single cell methylome, transcriptomic, and somatic genotyping reveals hypomethylation of PRC2 targets in *DNMT3A* R882 CH. **a,** UMAP dimensionality reduction (n = 528 cells) based on scRNA-seq data (Smart-seq2) after integration and batch correction of six plates (online methods). **b,** UMAP dimensionality reduction showing cluster gene markers for the transcriptome data. **c,** Number of CpG sites captured per cell after quality filtering (online methods). The metrics for each sample according to enzymatic digestion with Msp1 (Single) or Msp1 plus HaeIII (Double) are shown. **d,** Average single cell methylation at all regions (global, double digest), promoters, introns or exons. P-values from likelihood ratio test of LMM with/without mutation status (online methods). **e,** Average single cell methylation at CpH (i.e. CpA or CpT) sites. **f,** Average single cell methylation at 269 hypomethylated promoters identified with DMR analysis (shown in **Fig. 4e**, promoters with P-value < 0.05 and at least −5% methylation change) in CH02 and CH04. **g,** Average single cell methylation at SUZ12 (top panel) and EZH2 (bottom panel) ENCODE ChIP-seq peaks intersected with bivalently H3K27me3, H3K4me3-marked regions in CD34^+^ cells for CH02 and CH04. P-values from likelihood ratio test of LMM with/without mutation status. **h,** Normalized expression of PRC2 target genes with preferentially hypomethylated TSS (from **Fig. 4e**) in GoT data of WT versus MUT HSPCs. P-values from likelihood ratio test of LMM with/without mutation status. **i,** Comparison of average methylation values for TSS ± 1 kb regions in *DNMT3A* WT (n = 6) versus *DNMT3A* R882, *NPM1* mutated acute myeloid leukemia (AML; n = 7) samples in regions without (left) or with (right) PRC2 ChIP-seq peaks, controlling for CpG content. **j,** Comparison of average methylation values for promoter regions in WT (n = 122) versus *DNMT3A* R882 mutated AML (n = 9) samples from TCGA in regions without (left) or with (right) PRC2 ChIP-seq peaks, controlling for CpG content.

**Extended Data Figure 9.**
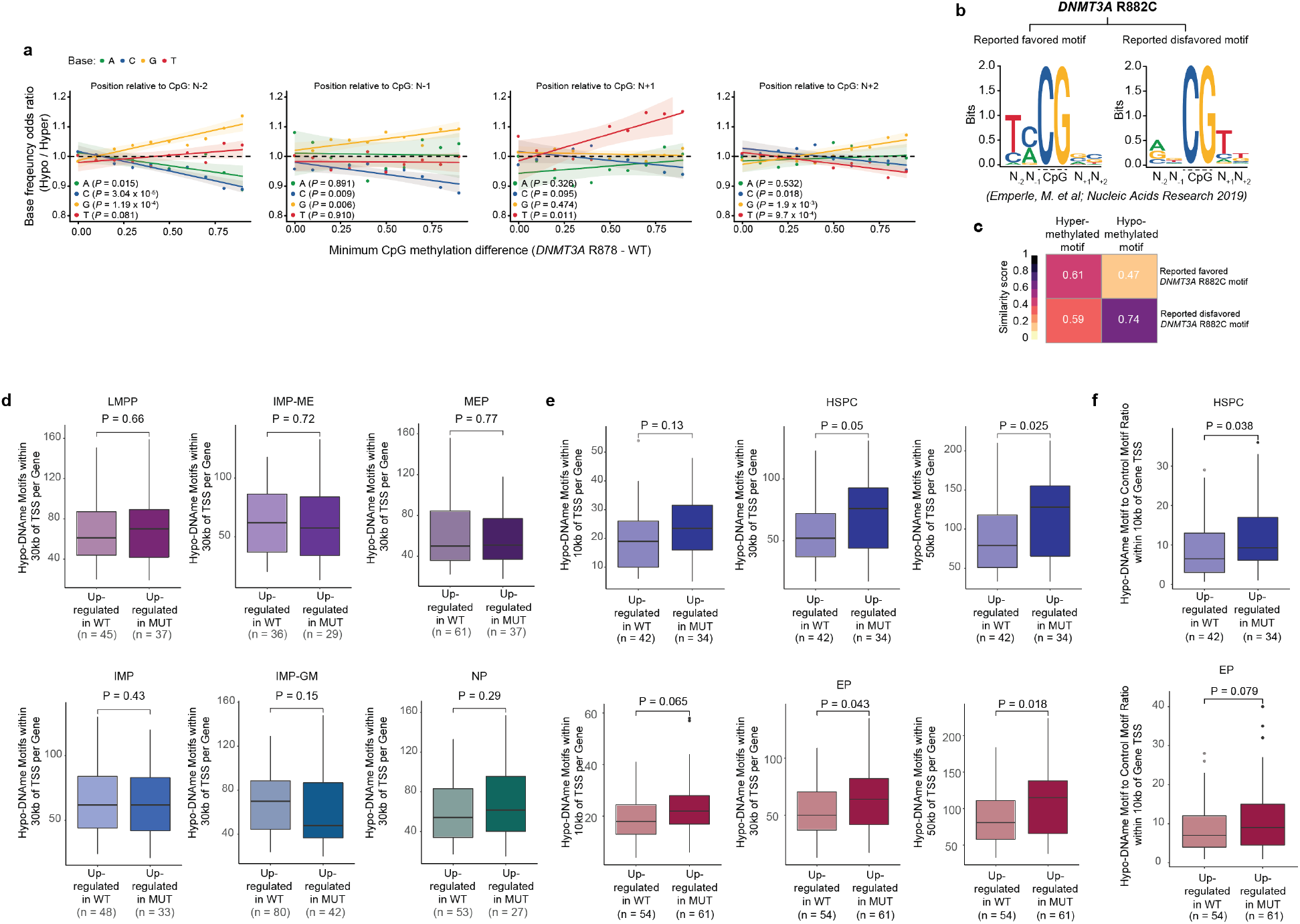
Motif enrichment at hypomethylated CpGs and hypomethylated motif enrichment in regions around differentially expressed genes. **a,** Base frequency odds ratio of hypo-versus hyper-methylated CpG flanking sequences at positions N-2, N-1, N+1, and N+2. The odds ratios were derived from base frequencies of flanking positions of the CpG sites hypo- or hyper-methylated in mutant versus wildtype cells above the thresholds shown in the x axis for minimum absolute CpG methylation difference (Pearson correlation, P-values derived from F-test). **b,** Reported motif logos derived from Emperle et al. for either hypomethylated (disfavored) or hypermethylated (favored) sites for DNMT3A R882 compared to its wildtype counterpart (left). **c,** Similarity scores between the reported and our de novo *DNMT3A* R882 hypo- and hypermethylated motifs as measured by correlation coefficients of the position weight matrices for the respective motifs <u>excluding</u> the CpG dinucleotide. **d,** Frequencies of *DNMT3A* R882 hypomethylated motif within 30kb of TSS of the differentially expressed genes between MUT and WT cells in progenitor subsets. P-values were calculated by Wilcoxon rank sum test. **e,** Frequencies of *DNMT3A* R882 hypomethylated motif within 10 kb, 30 kb or 50 kb of TSS of the differentially expressed genes between MUT and WT cells in HSPCs and EPs. P-values were calculated by Wilcoxon rank sum test. **f,** Ratio of frequencies of *DNMT3A* R882 hypomethylated motif to those of the control shuffled motif with CpG (**Fig. 5e**) within 10 kb of TSS of the differentially expressed genes between MUT and WT cells in HSPCs and EPs. P-values were calculated by Wilcoxon rank sum test.

**Extended Data Figure 10.**
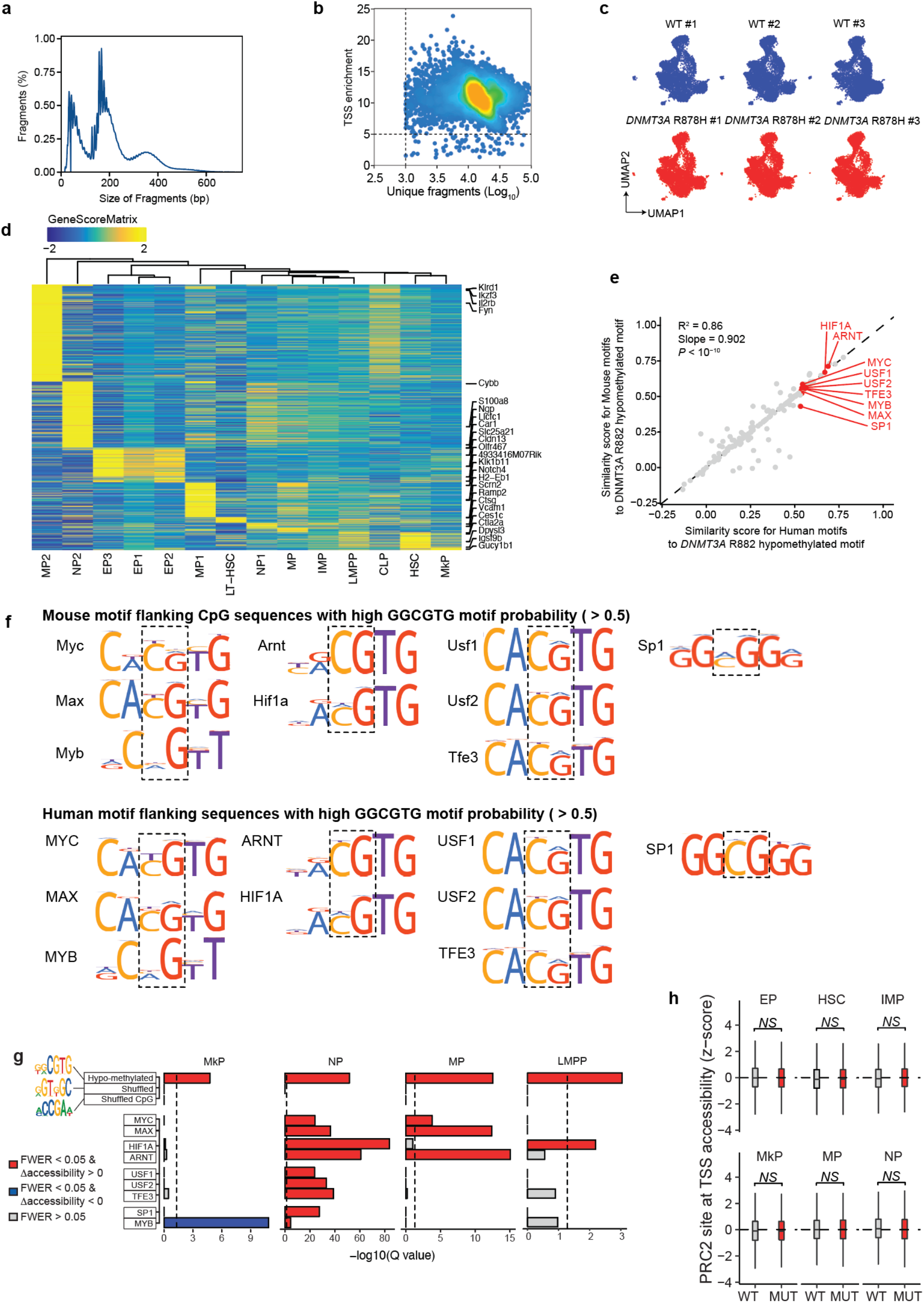
Single nucleus ATAC-seq of *Dnmt3a* R878H Lin-, c-Kit+ progenitors reveals enhanced accessibility of R882 hypomethylated motif and TF motifs with high similarity scores to the hypomethylated motif. **a,** Distribution of fragment size in snATAC-seq data of *Dnmt3a* R878H and wildtype Lin-, c-Kit+ progenitors (n = 3 in each cohort). **b,** TSS enrichment of accessible fragments as a function of unique fragments per cell. **c,** UMAP of integrated datasets *Dnmt3a* R878H and wildtype Lin-, c-Kit+ progenitors, displayed per sample (n = 3 in each cohort). **d,** Heatmap of gene accessibility scores for differentially accessible progenitor identity marker genes across progenitor subsets. **e,** Scatterplot of similarity scores of mouse TF motifs versus human TF motifs to the R882-hypomethylated motif (Pearson’s correlation, P-value derived from F-test). **f,** Binding motifs of mouse and human TFs with high similarity score to the R882-hypomethylated motif and expression in HSPCs (**Fig. 5b,** HOCOMOCO v11). **g,** FWER-adjusted P-values for accessibility changes between wildtype and *Dnmt3a* R878H cells by progenitor identities for hypo-methylated motif and shuffled motifs controls (with and without CpG), as well as motif accessibility deviation of the TFs identified **Fig. 5b (** related to **Fig. 5f)**. **h,** Accessibility of PRC2 targets between wildtype and *Dnmt3a* R878H and wildtype Lin-, c-Kit+ progenitor subsets.

**Extended Data Figure 11.**
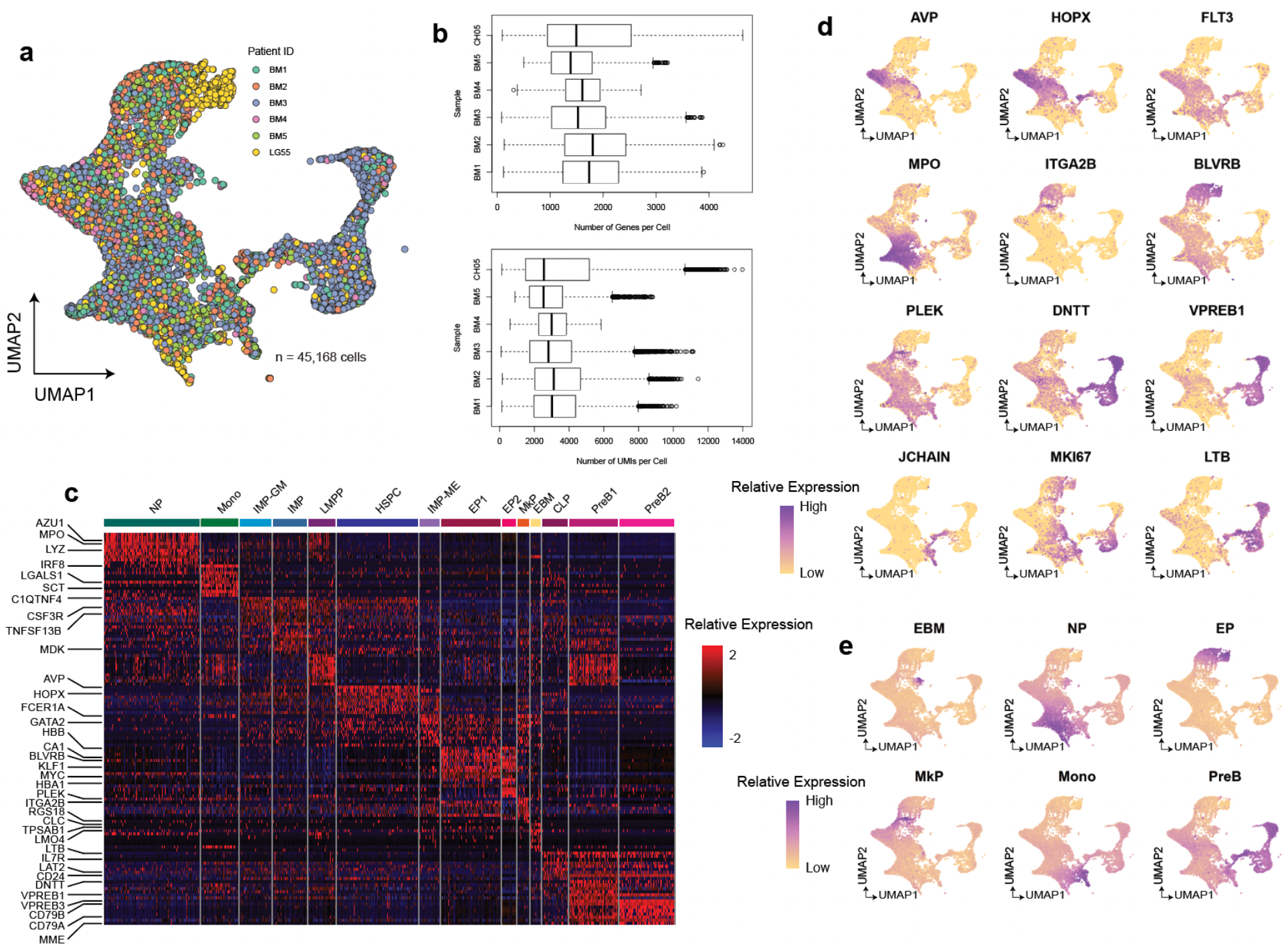
Integration of CH05 and control bone marrow CD34^+^ scRNA-seq data and assignment of progenitor subsets. **a,** UMAP of CD34^+^ progenitor cells from samples CH05 and samples BM01-05 after integration using the Seurat package (online methods). **b,** Number of genes per cell (top) and number of UMIs per cell (bottom) from CD34^+^ hematopoietic progenitors by patient sample after QC filters and down-sampling to equivalent geometric means of UMIs per patient. **c,** Heatmap of top 10 differentially expressed genes for progenitor subsets. **d,** UMAP representation of CD34^+^ cells showing cell marker gene expressions. **e,** Modules from Velten et al. (**Supplementary Table 2**) are scored and projected onto the UMAP representation of CD34^+^ cells.

**Extended Data Figure 12.**
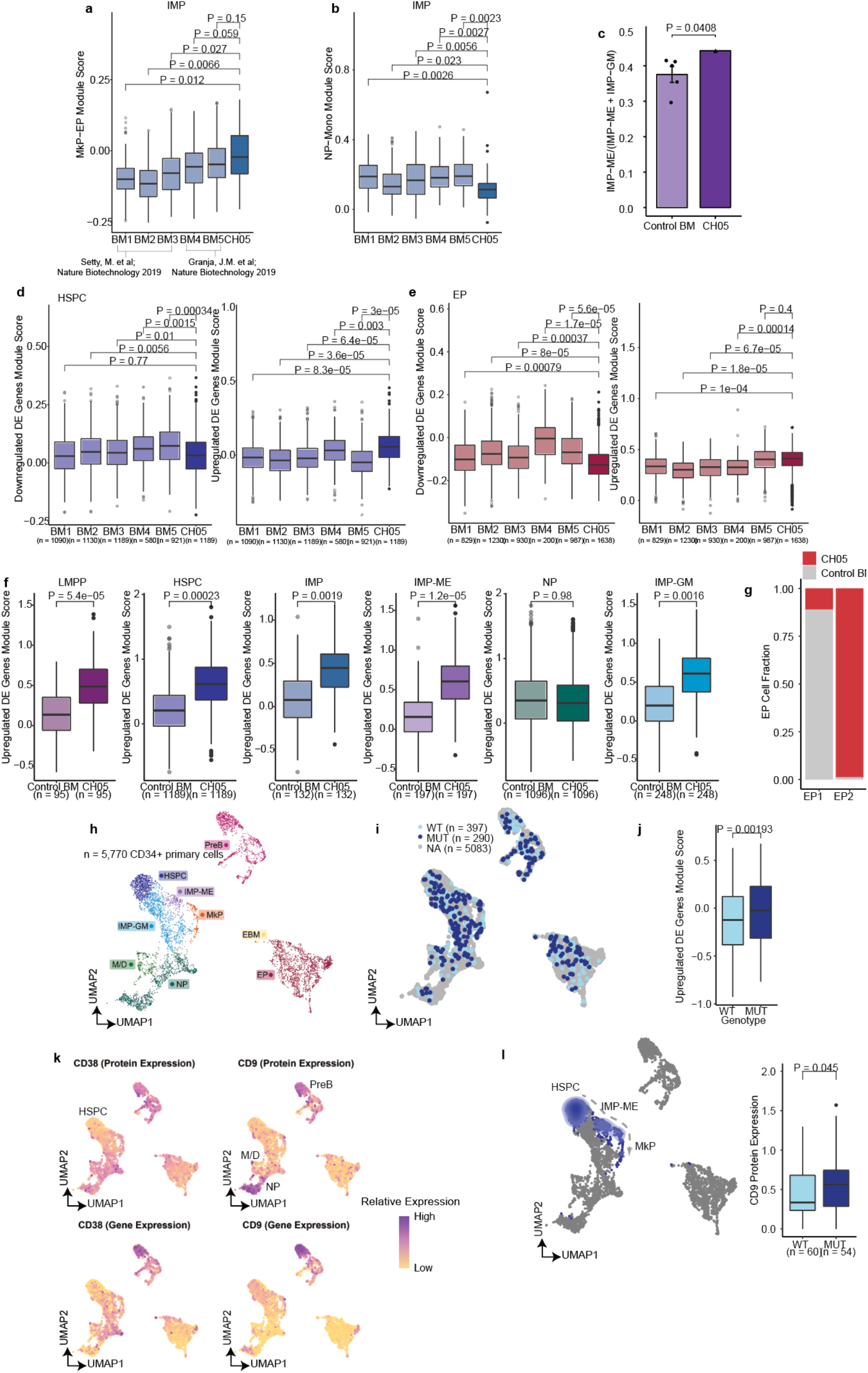
Bone marrow clonal hematopoiesis patient sample confirms results from CH01-CH04. **a,** Per-patient comparison of megakaryocytic-erythroid module scores in control bone marrow versus CH05 IMPs (**Supplementary Table 2**). Cell number downsampled to the same number (n = 132 cells per sample). P-values were calculated from likelihood ratio test of LMM with/without CH status. **b,** Per-patient comparison of granulocytic-monocytic module scores in control versus CH IMPs (**Supplementary Table 2**). P-values were calculated from likelihood ratio test of LMM with/without CH status. **c,** Fraction of IMP-ME cells out of all biased IMP (IMP-ME + IMP-GM) cells in control versus CH populations. P-value was calculated from one-sample t-test. **d,** Per-patient comparison of module scores for differentially down- or up-regulated genes in mutant *DNMT3A* HSPCs (identified in GoT data, **Fig. 3a,c**) in control versus CH HSPCs. P-values were calculated from likelihood ratio test of LMM with/without CH status. **e,** Per-patient comparison of module scores for differentially down- or up-regulated genes in mutant *DNMT3A* EPs (identified in GoT data, **Fig. 3a,c**) in control versus CH EPs. P-values were calculated from likelihood ratio test of LMM with/without CH status. **f,** Module scores for genes upregulated in at least 2 cell types (identified in GoT data, **Fig. 3b**) in control versus CH cells of major cell types. P-values from likelihood ratio test of LMM with/without CH status. **g,** Fraction of control BM or CH05 cells in EP1 versus EP2 cell clusters. **h**, UMAP of CH05 cells (clustered independently of the control BM samples) with progenitor cell assignments. **i**, UMAP of CH05 cells with genotyping data for WT (n = 397 cells) and *DNMT3A* R882 mutant (MUT; n = 290 cells). **j**, Normalized expression of differentially upregulated genes in at least 2 cell types, highlighted in **Fig. 3b** in wildtype versus mutated cells in CH05. **k**, UMAP of CH05 cells with protein expression (CITE-seq) and gene expression for CD38 and CD9. **l**, UMAP of CH05 cells highlighting HSPCs, IMP-ME, and MkPs (left) included in the comparison of CD9 expression in wildtype versus mutated cells (right).

**Extended Data Figure 13.**
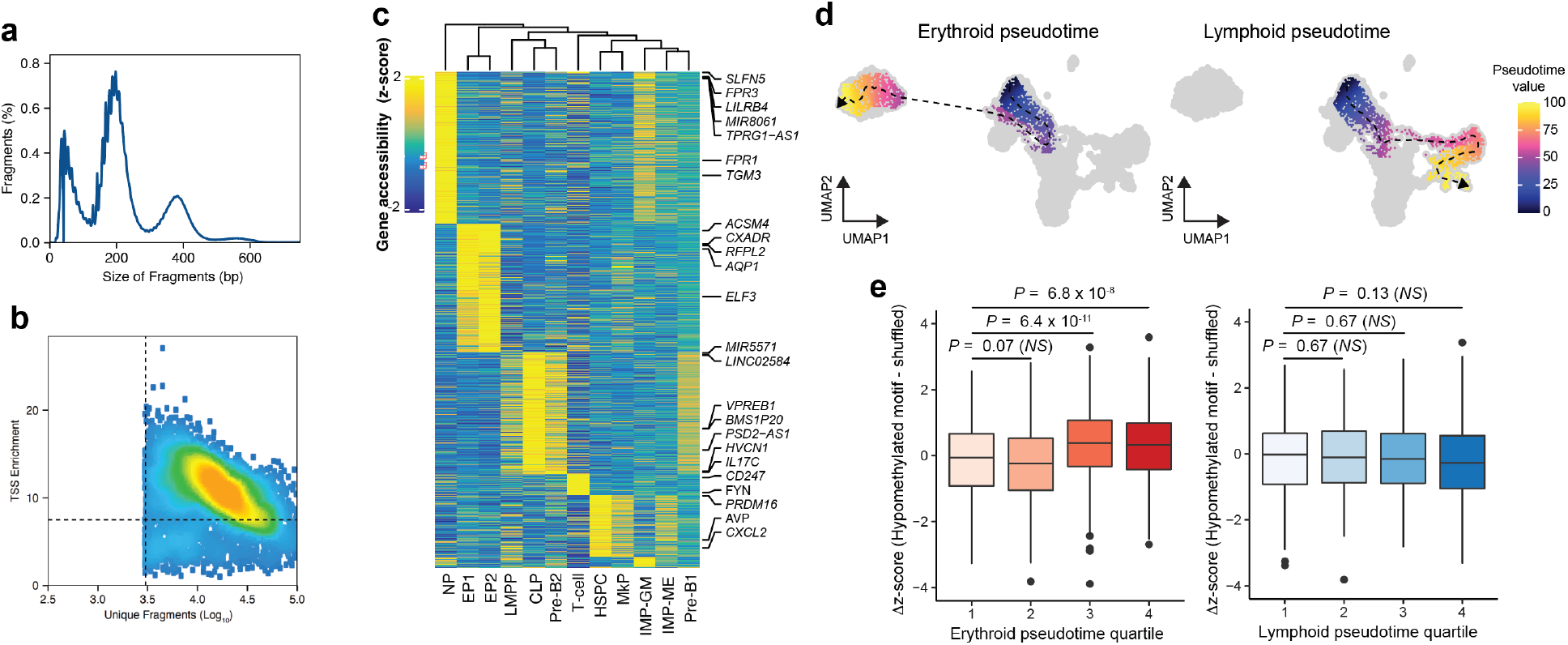
Single nucleus ATAC-seq data from bone marrow clonal hematopoiesis reveals enhanced accessibility of hypomethylated motif in mutated erythroid progenitors. **a,** Distribution of fragment size in snATAC-seq data of patient CH05 with *DNMT3A* R882 CH. **b,** TSS enrichment of accessible fragments as a function of unique fragments per cell. **c,** Heatmap of the gene accessibility scores for cluster marker genes (FDR < 0.01 and Log2FC > 1) by cell cluster. **d,** Pseudotime trajectories for either erythroid (left, n = 1,843 cells) or lymphoid (right, n = 1,740 cells) differentiation. **e,** Difference between hypomethylated and shuffled motif accessibility z-scores across either erythroid (n = 1,843 cells) or lymphoid (n = 1,740 cells) pseudotime trajectory quartiles. P-values were calculated by Wilcoxon rank sum test. HSPC, Hematopoietic stem and progenitor cell; IMP-ME, immature myeloid progenitor with megakaryocytic/erythroid bias; IMP-GM, immature myeloid progenitor with granulocyte/monocyte bias; LMPP, Lymphoid-myeloid pluripotent progenitor; MkP, Megakaryocyte progenitor; NP, Neutrophil progenitor; CLP, Common lymphoid progenitor; Pre-B1/2, Pre-B cell; EP1/2, Erythroid progenitor.

